# Engineering chimeric DNA polymerases for DNA movable type storage

**DOI:** 10.64898/2025.12.18.694503

**Authors:** Xutong Liu, Qixuan Zhao, Enyang Yu, Lijia Jia, Yue Shi, Di Liu, Haobo Han, Quanshun Li

**Affiliations:** Key Laboratory for Molecular Enzymology and Engineering of Ministry of Education, School of Life Sciences, Jilin University, Changchun 130012, China; State Key Laboratory of Virology and Biosafety, Wuhan Institute of Virology, Chinese Academy of Sciences, Wuhan 430071, China

**Author notes:** Corresponding authors. (Q. Li). (H. Han). (D. Liu).

**Keywords:** DNA-based information storage, DNA polymerase, DNA binding protein, processivity, DNA movable type storage

## Abstract

DNA-based information storage offers a promising alternative to conventional media due to its high density, long-term stability, and low energy requirements. However, its application remains hindered by synthesis costs, limited sequence length and poor scalability. DNA polymerase is a critical enzymatic tool in the DNA storage systems by enabling high-fidelity data writing and targeted sequence amplification. In this study, we engineered chimeric DNA polymerases by fusing the high-fidelity 9°N DNA polymerase with double-stranded DNA binding proteins derived from thermophilic archaea. These fusions significantly enhanced processivity, thermal stability, and salt tolerance by stabilizing enzyme-template interactions, mimicking sliding clamps while preserving catalytic efficiency. Leveraging these properties, we demonstrated precise file retrieval from a mixed oligonucleotide pool using orthogonal barcode primers. Compared with wild-type 9°N, the chimeric polymerases, particularly PLS, exhibited reduced substitution error rates and improved read accuracy. We then applied these enzymes to a DNA movable type storage system, where prefabricated DNA modules were assembled into encoding blocks. Using engineered polymerases, these blocks were recombined to enable flexible data rewriting without *de novo* DNA synthesis. Moreover, a multi-enzyme assembly strategy enabled the construction of kilobase-scale DNA sequences encoding a classical Chinese poem, achieving complete data recovery. All assembled fragments remained stable in *E. coli* over 100 generations, exhibiting the potential for *in vivo* storage. Collectively, our findings demonstrated the role of engineered DNA polymerases for DNA-based information storage. Moreover, this system reduced synthesis demands, supported scalable rewriting, and ensured long-term preservation, offering a practical route to sustainable, high-fidelity DNA data storage.

## 1. Introduction

Global digitization has driven explosive growth of digital data, creating an urgent demand for improved information storage media [**1,2**]. DNA, a natural genetic biomaterial, has emerged as a promising alternative to conventional digital storage media due to its high information density, long-term stability, and low energy demands during storage [**3–5**]. Currently, most DNA-based information storage utilizes “bit-to-base” encoding strategy, translating digital data into nucleotide sequences for chemical synthesis [**6–8**]. However, in practice, the synthesis of oligonucleotides is typically limited to lengths under 200 nt, constrained by cost, error rates and synthesis throughput [**9,10**]. As a result, storing large datasets usually requires a significantly higher number of oligonucleotides, which in turn increases the demand for extensive positional indexing to ensure accurate reconstruction from sequencing data [**11**]. This burden of positional indexing inevitably compromises the overall information density of storage system and contributes to higher synthesis cost. Moreover, each new file must be encoded into a newly synthesized set of DNA strands, leading to repeated synthesis even for structurally similar data [**12**]. Consequently, it is essential to develop storage strategies that minimize synthesis demands and support flexible and scalable information storage.

To reduce the cost and complexity of repeated DNA synthesis, DNA movable type (DNA-MT) storage encodes information using a fixed library of reusable, prefabricated DNA fragments. By selectively recombining these modular units, DNA-MT enables the dynamic construction of new sequences without additional synthesis, supporting scalable and versatile data storage [**13–16**]. For instance, Gong et al. constructed DNA-MT consisting of a 6-nt payload flanked by 43-nt primer arms and two Type IIS restriction enzyme sites, enabling ligation-based assembly [**13**]. Based on this concept, Wang et al. developed an automated inkjet printer that utilized pre-synthesized short oligonucleotides as DNA movable units [**14**]. The system leveraged enzymatic assembly of prefabricated units into larger constructs, supporting their repeated use and markedly reducing the need for *de novo* synthesis. Although these studies established conceptual foundation of DNA-MT, their practical applications remain limited by the inefficient assembly of short fragments into longer and complex DNA constructs. To address this challenge, various enzymatic assembly methods have been developed, such as polymerase cycling assembly (PCA) [**17**], Golden Gate assembly [**18**], and Gibson assembly [**19**]. Among these enzymatic strategies, DNA polymerases are indispensable for assembling DNA sequences, as they catalyze the extension of DNA strands with high specificity and fidelity. In our previous study, we demonstrated that the thermostable 9°N DNA polymerase, originally derived from *Thermococcus* sp. 9°N-7, could be used for encoding of diverse digital data types such as images, text, and videos into DNA, validating the feasibility of polymerase-mediated information writing [**20**]. Notably, 9°N DNA polymerase exhibited higher fidelity than commonly used commercial enzymes, which reduced base substitution errors during synthesis and improved the accuracy of data storage. However, 9°N and most *in vitro* DNA polymerases are non-replicative DNA polymerases and lack the accessory proteins, resulting in limited processivity and poor performance in long-fragment synthesis [**21,22**]. This limitation poses a particular challenge for DNA-MT storage system, which requires seamless and efficient assembly of multiple prefabricated DNA fragments into long constructs. Inspired by the *in vivo* DNA replication system, which employs accessory proteins such as sliding clamps to enhance polymerase-template association, we engineered 9°N DNA polymerase by fusing it with DNA binding proteins to improve processivity [**23–25**]. Among these candidate proteins, we selected the Sul7d family, derived from the thermophilic *Sulfolobus* species, due to their exceptional thermostability and strong non-specific sequence affinity for double-stranded DNA (dsDNA) [**26–30**]. These properties enable Sul7d to stably bind DNA under thermal cycling conditions, making it a promising fusion partner to enhance polymerase-template binding and sustain DNA strand synthesis.

In this study, we engineered chimeric DNA polymerases by fusing the 9°N DNA polymerase with archaeal DNA binding proteins, aiming to enhance processivity, thermal stability, and catalytic performance. These enzymes were then applied to DNA-based information storage, achieving precise file retrieval from mixed oligonucleotide pools, enzymatic assembly of modular DNA movable types, and construction of long-fragment DNA sequences for long-term *in vivo* storage. This enzyme-driven strategy reduced dependence on chemical synthesis and supported scalable, rewritable, and long-term *in vivo* DNA-based information storage systems.

## 2. Materials and methods

### 2.1 Materials

The oligonucleotide pools and all DNA sequences were chemically synthesized by GenScript (Nanjing, China). *E. coli* DH5α and *E. coli* BL21(DE3) were purchased from TransGen Biotech (Beijing, China). The pUC19 and pET-30a plasmids were maintained in our laboratory. Deoxyribonucleoside triphosphates (dNTPs), ssM13mp18 DNA, λDNA, BsaI-HFv2, and 10× ThermoPol buffer were purchased from New England Biolabs (Ipswich, MA). T4 DNA Ligase was purchased from TaKaRa Bio (Dalian, China). Ni-NTA agarose was purchased from Invitrogen (Waltham, MA). HiTrap Q HP column was purchased from GE Healthcare (Buckinghamshire, UK). Amicon centrifugal filter was purchased from Merck KGaA (Darmstadt, Germany). SanPrep Column DNA Gel Extraction Kit was purchased from Sangon Biotech (Shanghai, China). Hieff Clone Plus One Step Cloning Kit was purchased from Yeasen (Beijing, China).

### 2.1 Construction of chimerical DNA polymerases

Chimeric DNA polymerases were constructed by fusing the 9°N DNA polymerase with 7 kDa DNA binding proteins derived from *Sulfolobus* species *via* a flexible (GGGGS)_3_ linker. Each construct containing an N-terminal 6× His tag was cloned into the pET-30a plasmid between the NdeI and HindIII restriction sites. Specifically, PLS, PLA, and PLT polymerases incorporated the Sso7d (*Sulfolobus solfataricus*), Sac7d (*Sulfolobus acidocaldarius*), and Sto7d (*Sulfolobus tokodaii*) DNA binding proteins, respectively. The expression plasmids were synthesized and codon-optimized by GenScript (Nanjing, China). Related sequences of Sso7d, Sac7d and Sto7d were available in **Table S1**

### 2.2 Expression and purification of chimerical DNA polymerases

The plasmids were transformed into *E. coli* BL21(DE3) cells, which were cultured in LB medium at 37 °C until reach an OD_600_ of 0.6 to 0.8. Protein expression was induced by adding IPTG to a final concentration of 0.5 mM, followed by incubation for 4 h at 37 °C. Cells were collected by centrifugation, resuspended in lysis buffer (50 mM sodium phosphate buffer, 300 mM NaCl, 10 mM imidazole, 0.05% (v/v) Tween-20), and lysed *via* sonication. The lysate was heated to 85 °C for 20 min to denature endogenous proteins, followed by centrifugation at 12,000 × g for 30 min. The supernatant was incubated with Ni-NTA agarose at 4 °C for 1 h to bind His-tagged proteins. After washing with 20 mM imidazole buffer, the target protein was eluted with 75 mM imidazole. Imidazole was removed by Millipore ultrafiltration centrifuge tubes. Further purification was achieved *via* a HiTrap Q HP anion exchange column, and protein concentration was performed using a 30-kDa cutoff Amicon centrifugal filter. The purified DNA polymerase was verified by 10% SDS-PAGE and stored in buffer (10 mM Tris-HCl at pH 7.4, 100 mM KCl, 1 mM DTT, 0.1 mM EDTA, and 25% (v/v) glycerol) at -20 °C for further use.

### 2.3 Processivity assay of chimerical DNA polymerases

Processivity of DNA polymerase was assessed *via* a primer extension assay as previously described [**22, 31,32**]. In brief, 5’ FAM-labeled primer (M13FAM1) was added to ssM13mp18 DNA template in reaction buffer. This mixture was heated to 95 °C for 5 min, and then rapidly cooled on ice for at least 30 min to allow primer-template hybridization. DNA polymerases and dNTP were then added to initiate DNA synthesis at 72 °C. The reactions were terminated by the addition of 10 μL of 100 μM EDTA. Fragment analysis was performed to evaluate the processivity of each DNA polymerase by DNA sequencing. Electropherogram peak intensity were quantify to obtain the signal at each nucleotide position (nI) and the total peak intensity (nT) of all detectable products. The peaks of electropherogram traces were plotted and fitted to the following equation. The probability of not terminating at position I was calculated using the equation: PI = (nT-nI)/nT, where n represents the number of nucleotide residues incorporated and P_i_ is defined as the microscopic processivity at position I. The average primer extension length was determined using the formula 1/(1 ± PI). Additionally, the median length of the products of each reaction was determined as previously described by Wang et al. [**23**]. The PI and the average extension length were evaluated following analytical method proposed by Von Hippel et al. [**32**]. Related primer sequences were listed in **Table S2**.

### 2.4 Thermal stability of chimerical DNA polymerases

The purified DNA polymerases were incubated at 95 °C for varying durations to evaluate its thermal stability. Each PCR reaction mixtures contained 4 pM heated DNA polymerase for different time, 1 µM of each primer, 500 ng of pUC19 plasmid, 1× ThermoPol buffer, and 200 µM of each dNTPs. The PCR amplifications were performed with following protocol: initial denaturation at 95 °C for 30 s; 25 cycles of 95 °C for 30 s, 55 °C for 30 s, and 72 °C for 30 s; followed by a final extension at 72 °C for 5 min and a hold at 16 °C. PCR products from each heat treatment were analyzed by 1% agarose gel electrophoresis to evaluate the thermal stability of the DNA polymerases. The intensity of DNA band was quantified using Image J software (version 2.35, National Institutes of Health, Bethesda, MD). Related primer sequences were listed in **Table S2**.

### 2.5 Salt tolerance assay of chimerical DNA polymerases

The salt tolerance of DNA polymerases was evaluated under the same reaction conditions as described in “2.4 Thermal stability” section, except that the KCl concentration was adjusted to 0, 10, 20, 30, 40, 50, or 60 mM. PCR products were analyzed by agarose gel electrophoresis to assess the polymerase activity under different salt concentrations. The intensity of DNA band was quantified using Image J software (version 2.35, National Institutes of Health, Bethesda, MD). Related primer sequences were listed in **Table S2**.

### 2.6 PCR efficiency assay

To evaluate the amplification efficiency of each chimeric DNA polymerase, λDNA was used as the template. A series of primer pairs were designed to amplify fragments of 1, 2, 4, 6, and 8 kb in length. PCR reactions were performed under standard conditions, and the amplification efficiency of each polymerase was assessed by analyzing the yield and specificity of the resulting products *via* 1% agarose gel electrophoresis. Related primer sequences were listed in **Table S2**.

### 2.7 Encoding and decoding of data in oligonucleotide pools

Digital files, including “Jilin University” and “School of Life Sciences” logos, were individually encoded into DNA sequences using a Reed-Solomon coding to ensure error correction [**11**]. Each image was encoded into 5544 DNA strands with length of 102 nt. The inner code parameters were configured as: --N=34, --K=32, and --nuss=12, representing the length of inner code, message length, and number of symbols per sequence, respectively. According to file size, the outer code parameters were adjusted as: --n=1386, --k=1000, --numblocks=4 for the image, representing the length of outer code, original information, and number of blocks, respectively. The decoding process applied the same parameters as encoding.

### 2.8 Random access by DNA polymerases

For random access of data, each oligonucleotide was flanked by a pair of orthogonal primers corresponding to specific files. The designed constructs were listed as follows:

5’-TGGCTCATTTCACAATCGGT-[data]-TTGCACGGCAGGTCATTTAT-3’ (for Jilin University);

5’-ATAATTGGCTCCTGCTTGCA-[data]-TTGCACTTTCCGCCTACATT-3’ (for School of Life Sciences).

These oligo pools were synthesized using Customarray oligonucleotide chips by Genscript (Nanjing, China) and mixed in equal concentration (10 ng/μL) to form a combined file pool, totally containing 11088 DNA strands (each 143 nt). File-specific primer pairs (JLU-F/R or SLS-F/R) were used for selective PCR-based data access. PCR amplifications contained 10 ng template from mixed pool, 200 µM of each dNTP, 10 µM each primer, 1× ThermoPol buffer in a 100 μL volume. The thermal cycling protocol was set as described in “2.4 Thermal stability” section. The PCR products were purified by QIAquick PCR Purification Kit before sequencing. Related primer sequences were listed in **Table S2**.

### 2.9 Construction of DNA movable types

To ensure specific pairing, prevent unintended assembly, and enable downstream data operations, DNA movable types were classified into three categories: 20 nt address blocks (AMT) for positional identification, 20 nt checksum blocks (CMT) for error correction, and 50 nt payload blocks (PMT) for information encoding. A complete address unit consisted of three 20 nt sequences (An, Bn, and Cn). Each DNA movable type was flanked by a 5 nt linker to maintain assembly specificity and prevent unintended ligation. All DNA movable type sequences were designed using a computational scoring system based on the following criteria: The DNA movable types were designed by a computer-assisted method. A scoring system was developed to evaluate the specificity of these pairs: (1) no more than five consecutive A/T or four consecutive C/G bases; (2) C/G content between 20% and 80%; (3) inclusion of at least 3 types of bases; (4) no reverse complementary sequences longer than 4 nt; (5) no contiguous complementary sequences longer than 4 nt; (6) exclusion of pUC19 cloning sites, their reverse complementary sequences, and linker sequences; (7) a minimum Levenshtein distance of ≥ 7 for 20 nt DNA movable type or ≥ 15 for 50 nt DNA movable type. Linker sequences were selected from a pool of 256 randomly generated 5 nt sequences and screened by following criteria: (1) inclusion at least two different types of bases; (2) no more than three consecutive identical bases; (3) GC content between 20% and 80%. The corresponding sequences of each character in Chinese poem “wang lu shan pu bu” was listed in **Table S3-S5**.

### 2.10 Assembly of DNA movable type blocks (DMTBs) *via* overlap PCR

To assemble complete DNA movable type blocks (DMTBs), AMT, CMT, and PMT movable type units were enzymatically assembled using overlap extension PCR. Each 50 μL reaction mixture contained 8.0 pM of PLS DNA polymerase, 1 μM of each DNA movable types, 10 mM dNTPs, and 5μL of 10× ThermoPol buffer. The PCR conditions were as follows: initial denaturation at 95 °C for 5 min; 25 cycles of 95 °C for 30 s, 55 °C for 30 s and 72 °C for 30 s; and a final extension at 72 °C for 10 min. The PCR products were purified using Gel Extraction Kit to remove non-specific products. The assembled DNA constructs were subsequently inserted into pUC19 plasmid *via* homologous recombination using Hieff Clone Plus One Step Cloning Kit and then transformed into *E. coli* DH5α for *in vivo* storage.

### 2.11 Assembly of DNA movable types *via* PCA

To construct a continuous DNA fragment from DNA movable types, eight oligonucleotides were chemically synthesized. These oligonucleotides were designed to possess 20 bp overlapping regions with adjacent sequences to facilitate assembly *via* polymerase cycling assembly (PCA). Specifically, PCA-1 and PCA-8 served as outer primers, while PCA-2 through PCA-7 were internal fragments. The PCR reaction mixture contained 1 µM of each internal oligonucleotide (PCA-2 to PCA-7), 100 µM of each outer oligonucleotide (PCA-1 and PCA-8), 10 mM dNTPs, 1× ThermoPol buffer, and nuclease-free water to a final volume of 50 µL. PLS DNA polymerase was added at a final concentration of 4 pM. Thermal cycling conditions were as follows: initial denaturation at 95 °C for 5 min; 25 cycles of 95 °C for 30 s, 55 °C for 30 s, and 72 °C for 30 s; with a final extension step at 72 °C for 10 min. The full-length product was verified by 1% agarose gel electrophoresis and purified using the Gel Extraction Kit. Related primer sequences were listed in **Table S2**.

### 2.12 Assembly of DNA movable type units (DMTUs) and DMT-Files (DMTFs)

DMTUs and DMTFs were constructed by assembling pre-formed DNA movable type blocks (DMTBs) using the strategy of Golden gate assembly and Gibson assembly. Briefly, each DMTB fragment was amplified from the engineered pU19 plasmid using PLS DNA polymerase. BsaI recognition sites were introduced at both ends during primer design to enable seamless ligation. For digestion, 300 ng of PCR products was mixed with 5 μL of 10× buffer, and 1 μL of BsaI-HFv2 restriction enzyme, followed by incubation at 37 °C for 1 h. The digested DNA fragments were purified using the Gel Extraction Kit before assembly. For the ligation, 300 ng of purified DNA fragments were combined with 2 μL of 10× buffer, and 2 μL of T4 DNA ligase, and ddH_2_O was added to a final volume of 20 μL. The mixture was incubated at 4 °C overnight for DNA ligation. The resulting DMTUs and DMTFs were verified by 1% agarose gel electrophoresis and subsequently transformed into E. coli DH5α for *in vivo* storage.

### 2.13 Data decoding of DNA movable type storage

Sanger sequencing results were decoded using predefined linker sequences and the known lengths of each DNA movable type. Regular expressions were used to parse the sequences into the structure “AMT (An-Bn-Cn)-CMT-PMT”, allowing for the identification of individual movable type components. Each extracted sequence was compared to entries in a reference codebook using fuzzy string matching. The algorithm calculated sequence similarity and assigned the corresponding movable type ID to the best-matching reference. Sequences with a similarity score below 85% were considered unmatched and labeled as “NA.” Checksum blocks were then used to validate each decoded sequence. If the checksum did not match the expected value, the entire movable type combination was identified as an error. For sequences passing checksum verification, the address (An-Bn-Cn) was reconstructed to determine the correct position of the payload. The payload blocks were subsequently arranged in order, and each movable type ID was mapped to its corresponding character using the codebook, thereby reconstructing the original encoded information.

## 3. Results and Discussion

### 3.1 Expression and characterization of chimerical DNA polymerases

To enhance the processivity of non-replicative DNA polymerases, we constructed chimeric enzymes by genetically fusing the 9°N DNA polymerase with dsDNA binding proteins. As DNA binding proteins, we selected Sso7d, Sto7d, and Sac7d, small and thermostable proteins derived from the hyperthermophilic archaea *Sulfolobus solfataricus*, *Sulfolobus tokodaii*, and *Sulfolobus acidocaldarius*, respectively [**33-35**]. These proteins are known for their non-sequence-specific binding to dsDNA and their role in maintaining genome integrity under extreme environmental stress, making them suitable candidates for enhancing DNA polymerase-template association. To generate the chimeric constructs, each DNA binding domain was fused to the C-terminus of 9°N DNA polymerase *via* a flexible peptide linker (GGGGS)_3_, which was introduced to reduce the steric hindrance and maintain the independent folding of each domain (**Fig. 1A** and **1B**). The resulting fusion enzymes were designated as PLS, PLT, and PLA, corresponding to the incorporation of Sso7d, Sto7d, or Sac7d, respectively.

**Fig. 1.**
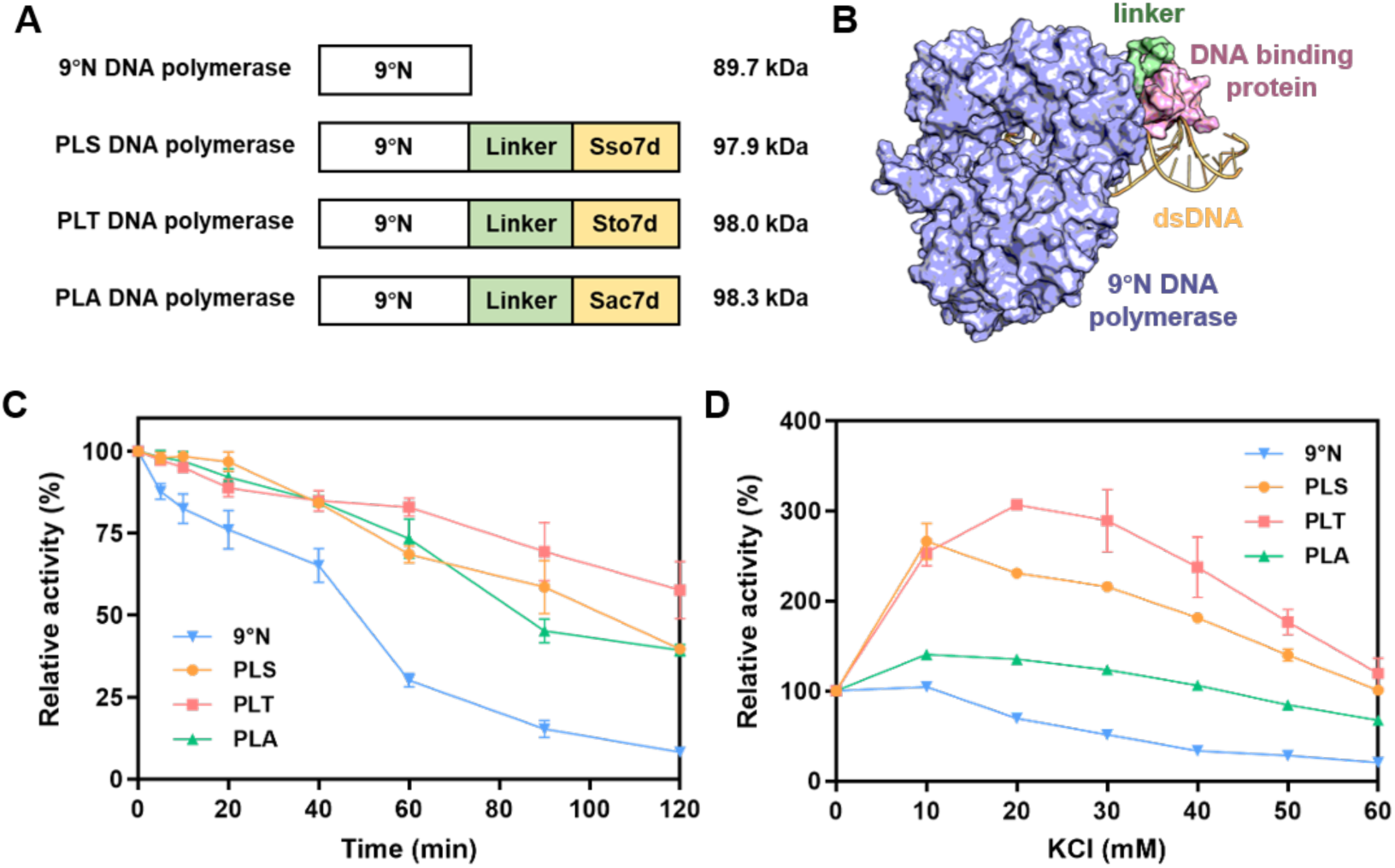
Structural and biochemical characterization of 9°N DNA polymerase and chimeric variants. (**A**) Schematic domain architecture of 9°N and chimeric DNA polymerases. (**B**) Structural modeling of chimeric DNA polymerases PLS. (**C**) Thermal stability analysis of 9°N and chimeric DNA polymerases. (**D**) Salt tolerance analysis of 9°N and chimeric DNA polymerases.

The genes encoding PLS, PLT, and PLA DNA polymerases were individually cloned into the pET-30a vector and transformed into *E. coli* BL21(DE3) cells. Protein expression was induced by IPTG, and the recombinant proteins were subsequently purified using nickel affinity chromatography, followed by cation exchange chromatography. SDS-PAGE analysis showed a single band at approximately 89 kDa for 9°N DNA polymerase, and at approximately 98 kDa for each chimeric DNA polymerases, consistent with their theoretical molecular weights predicted by Expasy based on their amino acid sequences (**Fig. S1**). Since archaeal DNA binding proteins contribute to genomic stability under extreme conditions, we next investigated whether their fusion enhanced the stability of DNA polymerase. Thermal stability, which was a critical characteristic for PCR applications, was assessed by pre-incubating the enzymes at 95 °C for different time periods, followed by PCR analysis. As shown in **Fig. 1C and Fig. S2**, the chimeric DNA polymerases PLS, PLT, and PLA exhibited significantly higher catalytic activity than 9°N after heat treatment. Notably, after incubation at 95 °C for 90 min, the 9°N DNA polymerase retained less than 15% of its activity, whereas PLS, PLT, and PLA retained 58%, 69%, and 45% of their initial activities, respectively. These results indicated that fusion with archaeal DNA binding proteins substantially improved the thermal stability of DNA polymerases. To further evaluate the robustness of the chimeric enzymes under other challenging conditions, we examined their salt tolerance by testing the performance in increasing concentrations of KCl. As shown in **Fig. 1D** and **S3**, the catalytic activity of 9°N DNA polymerase decreased to 69% in the presence of 20 mM KCl, indicating that even low concentrations of salt could substantially inhibited its enzymatic activity, likely by disrupting polymerase-template interactions. In contrast, the chimeric polymerases PLS, PLT, and PLA maintained higher catalytic activities of 230%, 306% and 135%, respectively, under the same conditions, which were even higher than those observed in salt-free environments. This phenomenon suggested that fusion with DNA binding domains significantly enhanced the salt tolerance of DNA polymerases, enabling the chimeric enzymes to maintain high catalytic activity even under ionic stress. In particular, low concentrations of KCl further promoted polymerase-template interactions of chimeric enzymes, likely by stabilizing electrostatic contacts and reducing repulsive forces. However, as the KCl concentration increased, the activity of all three chimeric enzymes gradually declined. This reduction was likely due to disruption of electrostatic interactions and conformational destabilization, which in turn impaired template binding and reduced catalytic efficiency. To further investigate the catalytic performance of the chimeric DNA polymerases, we assessed their processivity using a fluorescence-based primer extension assay. Specifically, microscopic processivity (PI) was determined by measuring the length of the extended DNA fragments and their corresponding fluorescence signal intensities. The average primer extension length was then calculated based on the PI values, serving as an indicator of polymerase processivity. As shown in **Table 1**, 9°N DNA polymerase yielded an average primer extension length of 13.7 nt. In contrast, chimeric enzymes PLS, PLT, and PLA achieved significantly longer extension length of 26.8 nt, 29.7 nt, and 22.9 nt, respectively. These results indicated that fusion with DNA binding domains significantly enhanced the polymerase processivity, supporting DNA binding proteins fusion as an effective strategy for improving polymerase performance.

**Table 1.** Processivity analysis of 9°N and chimeric DNA polymerases.

| Enzymes | Microscopic processivity<br>(PI) | Average primer extension length<br>(nt) [1/(1 ± PI)] |
| --- | --- | --- |
| 9°N | 0.927 ± 0.01 | 13.7 ± 2.1 |
| PLS | 0.962 ± 0.003 | 26.8 ± 1.7 |
| PLT | 0.966 ± 0.004 | 29.7 ± 3.4 |
| PLA | 0.956 ± 0.001 | 22.9 ± 0.3 |

Encouraged by the enhanced processivity of the chimeric polymerases, we next evaluated their PCR efficiency in amplifying DNA fragments of increasing lengths. As shown in **Fig. 2A**, 9°N DNA polymerase exhibited limited amplification capability, producing a clear band at 1 kb and only partial amplification at 2 kb under a 30 s extension time. In contrast, the chimeric DNA polymerases showed markedly enhanced performance under the same conditions. Specifically, both PLT and PLA successfully amplified a 6 kb fragments, while PLS yielded a 6 kb product and showed a partial amplification of 8 kb fragment, indicating superior strand extension capability and greater tolerance to salt inhibition. When the extension time was increased to 1 min at 72 °C, similar results were observed (**Fig. 2B**). 9°N DNA polymerase was able to amplify a 2 kb fragment, whereas the chimeric polymerases PLS, PLT, and PLA successfully amplified fragments up to 8 kb. These findings reinforced the conclusion that the fusion of DNA binding proteins could enhance the long-fragment amplification of polymerase. To assess the impact of salt, we compared amplification performance under salt-free and 20 mM KCl conditions (**Fig. 2** and **S4**). For 9°N DNA polymerase, the presence of salt reduced its amplification capacity from 4 kb to 2 kb under a 30 s extension time, indicating that salt ions inhibited its catalytic activity (**Fig. S4A**). In contrast, the chimeric polymerases showed enhanced performance in the presence of KCl, achieving 8 kb amplification compared to 6 kb under salt-free condition. This enhancement highlighted the improved salt tolerance conferred by the fused DNA binding proteins, which stabilized polymerase-template interaction and enabled more efficient amplification under salt conditions.

**Fig. 2.**
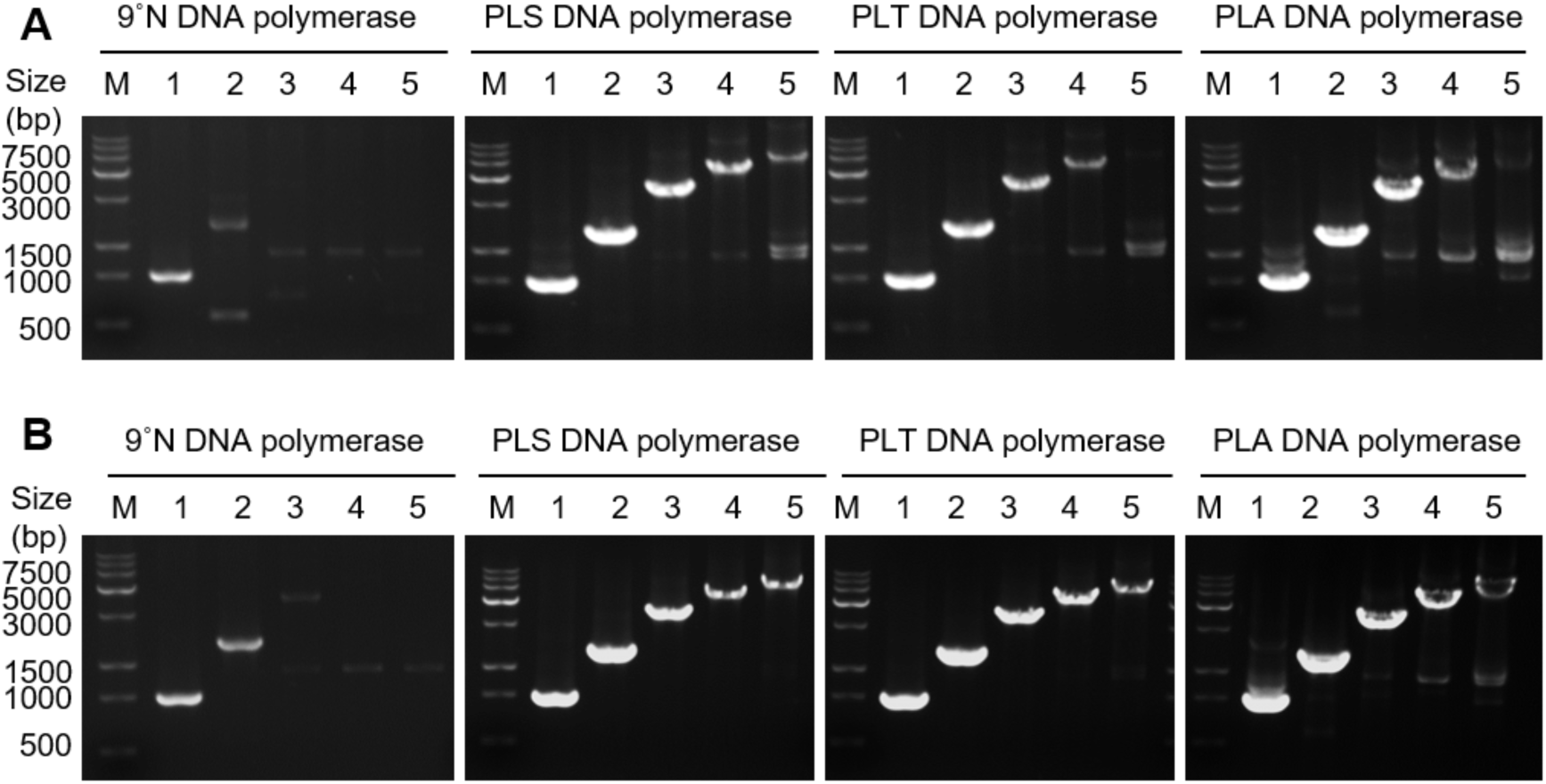
Evaluation of PCR efficiency using λDNA as template under different thermocycling conditions in the presence of 20 mM KCl. (**A**) PCR with 30 s extension at 72 °C; (**B**) PCR with 60 s extension at 72 °C. M: Marker; Lanes 1-5: target amplicons of 1, 2, 4, 6, and 8 kb, respectively.

### 3.2 Molecular insights into the increased processivity of chimeric DNA polymerases

To investigate the molecular insights into the increased processivity of chimeric DNA polymerases, we firstly performed sequence alignment and secondary structure prediction of three fused DNA binding proteins. As shown in **Fig. S5**, sequence comparison revealed that Sso7d shared 91% identity to Sto7d, and 81% identity to Sac7d, indicating high conservation among these homologs. Most amino acid residues were conserved across the three proteins, with sequence variations mainly occurring at the N- and C-terminal regions. Moreover, secondary structure prediction showed strong structural similarity among the three DNA binding proteins, with only minor differences observed in loop regions and their adjacent structures (**Fig. S6**). These results indicated that the three DNA binding proteins shared a conserved structural framework, making them suitable for fusion with 9°N DNA polymerase without disrupting protein folding or enzymatic function. The minor structural differences were unliked to interfere with overall protein folding, suggesting that any functional variations among the chimeric enzymes were more likely due to subtle distinctions in protein-DNA interaction rather than major structural incompatibility.

Based on the structural analysis, we next performed 500 ns Gaussian accelerated molecular dynamics (GaMD) simulations [**36**] on 9°N, PLS, PLT, and PLA polymerases in complex with dsDNA to evaluate their dynamic behavior of polymerase/DNA interaction. During the simulations, all chimeric DNA polymerase/DNA complexes exhibited an initial increase in root-mean-square deviation (RMSD), followed by stabilization between 15 Å and 20 Å (**Fig. S7)**. This convergence indicated that each complex reached a relatively stable conformation over time. Root-mean-square fluctuation (RMSF) analysis was performed to evaluate the flexibility of residues in the chimeric DNA polymerases throughout the simulation trajectories (**Fig. S8**). Notably, the highest fluctuations were observed in the DNA binding domains of the chimeric polymerases, which likely resulted from their swinging state during the initial dynamic association with dsDNA to achieve stable binding. The radius of gyration (Rg) was used to evaluate the structural compactness of the chimeric polymerase-DNA complexes, with smaller Rg values indicating more compact protein conformations (**Fig. S9**). Compared to the 9°N DNA polymerase, all three chimeric DNA polymerases exhibited reduced Rg values, suggesting that the fusion of DNA binding domains enhanced the overall compactness, likely due to strengthened interactions between DNA polymerase and DNA binding protein.

The binding free energies between the chimeric DNA polymerases and dsDNA were calculated using molecular mechanics Poisson Boltzmann surface area (MM-PBSA) [**37**] and molecular mechanics Generalized Born and Surface Area (MM-GBSA) [**38**]. As shown in **Table 2**, the 9°N DNA polymerase exhibited the least negative binding energy among four enzymes, indicating relatively weak binding to dsDNA. In contrast, PLS, PLT and PLA showed substantially more negative binding energy, suggesting stronger and more stable interaction with dsDNA. These findings revealed that the fused DNA binding domains significantly enhanced polymerase-DNA affinity, which contributed to the improved processivity. Notably, among these chimeric enzymes, PLS demonstrated both strong dsDNA binding and superior long-fragment amplification capability, which was consistent with its high processivity performance. Interestingly, although PLA DNA polymerase exhibited a more negative binding energy than the PLS, its long-fragment amplification ability was inferior. This result indicated that excessively strong DNA binding might hinder the polymerase dissociation and translocation, ultimately inhibiting processivity. To further evaluate polymerase-DNA interaction, we analyzed the surface electrostatic potential of DNA binding domains (**Fig. S10**). The DNA binding interfaces of all three domains exhibited predominantly positive charge, promoting interactions with the negatively charged phosphate backbone of dsDNA. Notably, the three DNA binding domains exhibited differences in both the distribution and charge strength of positive electrostatic potentials at their contact sites with DNA.

**Table 2.** Free binding energy of 9°N and chimeric DNA polymerases.

| Enzymes | MM-PBSA | MM-GBSA |
| --- | --- | --- |
| | $\Delta G_{\text{binding}}$ (kcal/mol) | $\Delta G_{\text{binding}}$ (kcal/mol) |
| 9°N | -41.4754 | -223.6795 |
| PLS | -109.4642 | -270.5023 |
| PLT | -74.8216 | -252.1683 |
| PLA | -1417.4042 | -1581.3059 |

To investigate the structural determinants underlying the enhanced processivity, we performed contact fraction analysis of the MD trajectories to identify the specific interaction between key residues of the DNA binding domains and dsDNA (**Fig. 3**). This analysis identified specific interactions between substrate nucleotides and DNA binding domains of chimeric DNA polymerase, including hydrogen bonds, π-π stacking, salt bridges, and electrostatic interactions. The three DNA binding domains exhibited similar interaction patterns with dsDNA, likely due to their sequence homology. Among these, Trp24 in the Sso7d domain of the PLS was identified as a critical residue for π-π stacking with dsDNA binding, forming two stable interactions at distance of 4.6 Å and 4.7 Å, respectively (**Fig. 3A**). The critical importance of Trp24 in DNA binding was further supported by previous findings, where its mutation led to a marked reduction in polymerase processivity [**23**]. In Sto7d domain of PLT, Arg25 formed a stable hydrogen bond with dsDNA at a distance of 1.8 Å (**Fig. 3B**). In Sac7d domain of PLA, Lys7, Tyr8, and Lys9 formed hydrogen bonds with dsDNA at distances of 1.9 Å, 2.1 Å, and 2.9 Å, respectively (**Fig. 3C**). These extensive polar interactions might account for the higher binding free energy of PLA, as observed in binding energy analysis. Notably, all three DNA binding proteins predominantly interacted with the DNA minor groove of dsDNA, functioning similarly to a sliding clamp to anchor the chimeric DNA polymerase to DNA and improve binding stability and processivity. While the overall binding mode was conserved across the three domains, subtle differences in interaction types, such as hydrogen bonds and π-π stacking interactions, likely contributed to the variations in processivity among chimeric enzymes. Collectively, these findings provided a comprehensive understanding into how DNA binding proteins enhanced the processivity and stability of chimeric DNA polymerases, supporting further engineering of high-performance DNA polymerases.

**Fig. 3.**
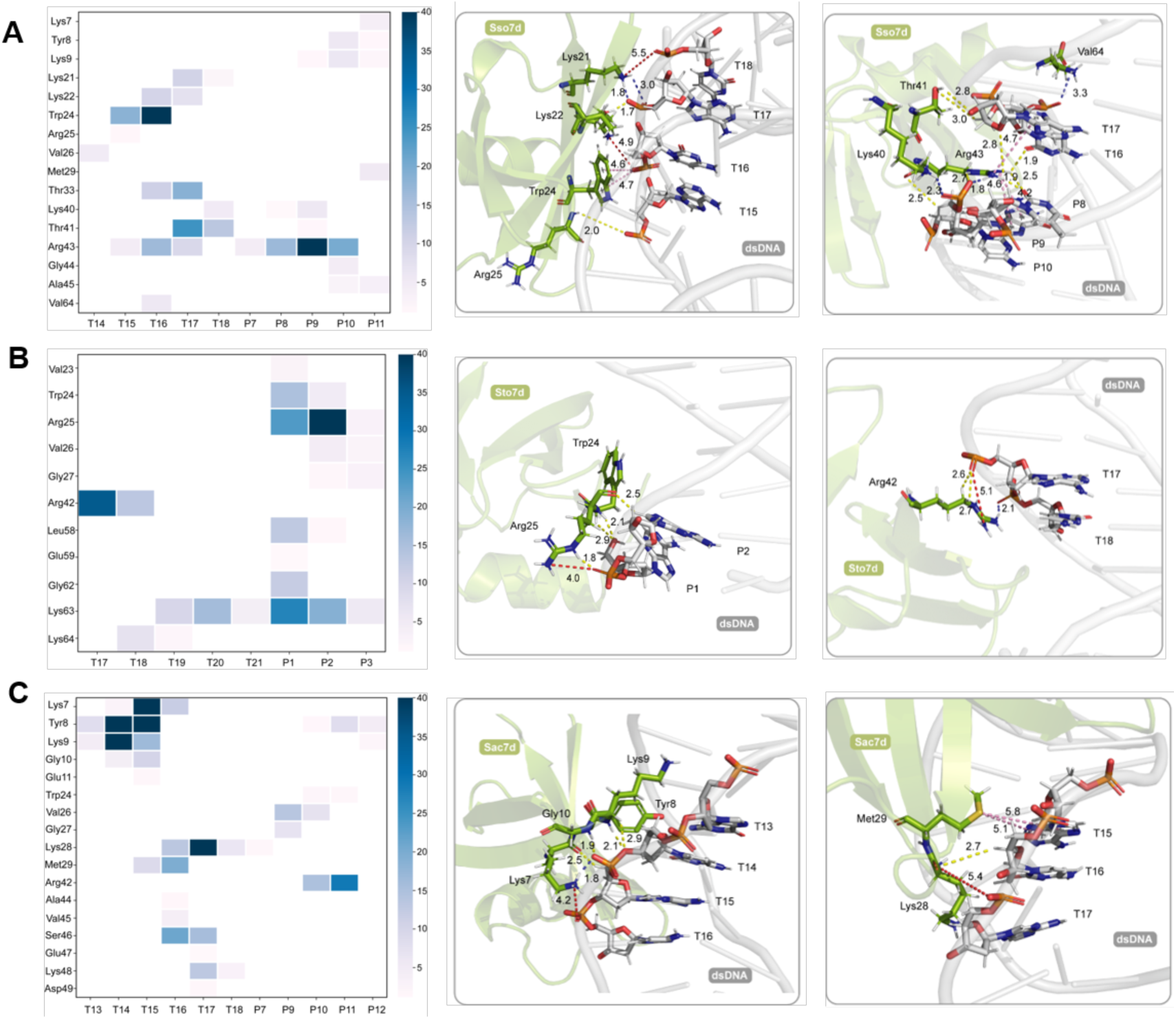
Interaction patterns of chimeric DNA polymerases with dsDNA. Molecular dynamics simulations of interactions between (**A**) PLS, (**B**) PLT, and (**C**) PLA DNA polymerases and dsDNA by contact fraction analysis and structural mapping of interacting residues. Hydrogen bonds were shown in yellow, π-π stacking in pink, charge-charge interactions in red, and salt bridges in purple.

### 3.3 Chimeric DNA polymerase enabled file retrieval in DNA-based information storage

Random data access is crucial requirement for practical DNA-based information storage, enabling the efficient and reliable information retrieval of specific files without sequencing the entire dataset. PCR-based strategies are commonly employed for data access in DNA information storage, as they rely on the primer-template complementarity to selectively amplify target barcode sequences for accurate file retrieval. To enhance the accuracy and robustness of data retrieval, we developed an orthogonal primer-based access system (**Fig. 4A**). To test the performance of the system in multi-file data retrieval, we encoded and stored two representative image files: the logo of Jilin University and the logo of the School of Life Sciences at Jilin University. To improve image fidelity and increase the complexity of addressing, color normalization was applied during encoding. Each data files were individually converted into an oligonucleotide pool using Reed-Solomon coding [**11**], generating two sets of 5,544 oligonucleotides, each 102 nucleotides in length. Orthogonal barcode sequences were appended to both ends of each sequence to enable file retrieval. The two oligonucleotide pools were then physically mixed at equal concentrations to evaluate the specificity of file retrieval in a mixed oligo pool. Specifically, the file encoding Jilin University logo was selectively retrieved using its corresponding orthogonal primers. PCR was performed with each DNA polymerases, followed by high-throughput sequencing and Reed-Solomon-based decoding of retrieved sequences. As shown in **Fig. 4B**, all four polymerases successfully amplified the desired file, demonstrating the efficiency of the orthogonal primer system for targeted file access. Moreover, compared with the 9 ° N DNA polymerase, the three chimeric DNA polymerases exhibited higher fidelity during amplification, resulting in fewer sequence errors. This improvement indicated that DNA binding proteins improved the fidelity of the chimeric DNA polymerases, promoting accurate DNA synthesis. Furthermore, they reduced the incidence of common amplification errors such as deletions, insertions, and substitutions, highlighting their potential to enhance the accuracy and the reliability of random addressing in DNA-based information storage. Since substitution errors in DNA information storage are primarily introduced during PCR amplification, the fidelity of DNA polymerases directly influences the accuracy of data retrieval. To further investigate amplification fidelity, substitution matrix analysis was performed to quantify the misincorporation during file retrieval (**Fig. 4C-4F**). Compared to the 9 ° N DNA polymerase, all three chimeric DNA polymerases significantly reduced substitution errors, including transitions and transversions, during template amplification. For instance, when the thymine (T) was expected base, the total substitution error rates was 3.02 × 10^-4^ for 9° N DNA polymerase, while the rates were reduced to 1.18 × 10^-4^ for PLS, 2.00 × 10^-4^ for PLT, and 2.24 × 10^-4^, respectively. These results demonstrated that the chimeric DNA polymerases exhibited improved fidelity in both DNA amplification and random access addressing. Specifically, PLS DNA polymerase achieved low substitution error rates and produced a high proportion of correct reads, which accounted for 98.79% of all sequenced reads. This enhanced fidelity reduced non-specific amplification and improved the accuracy of targeted file retrieval, supporting the application of chimeric DNA polymerases for precise and reliable data access in future large-scale DNA storage systems.

**Fig. 4.**
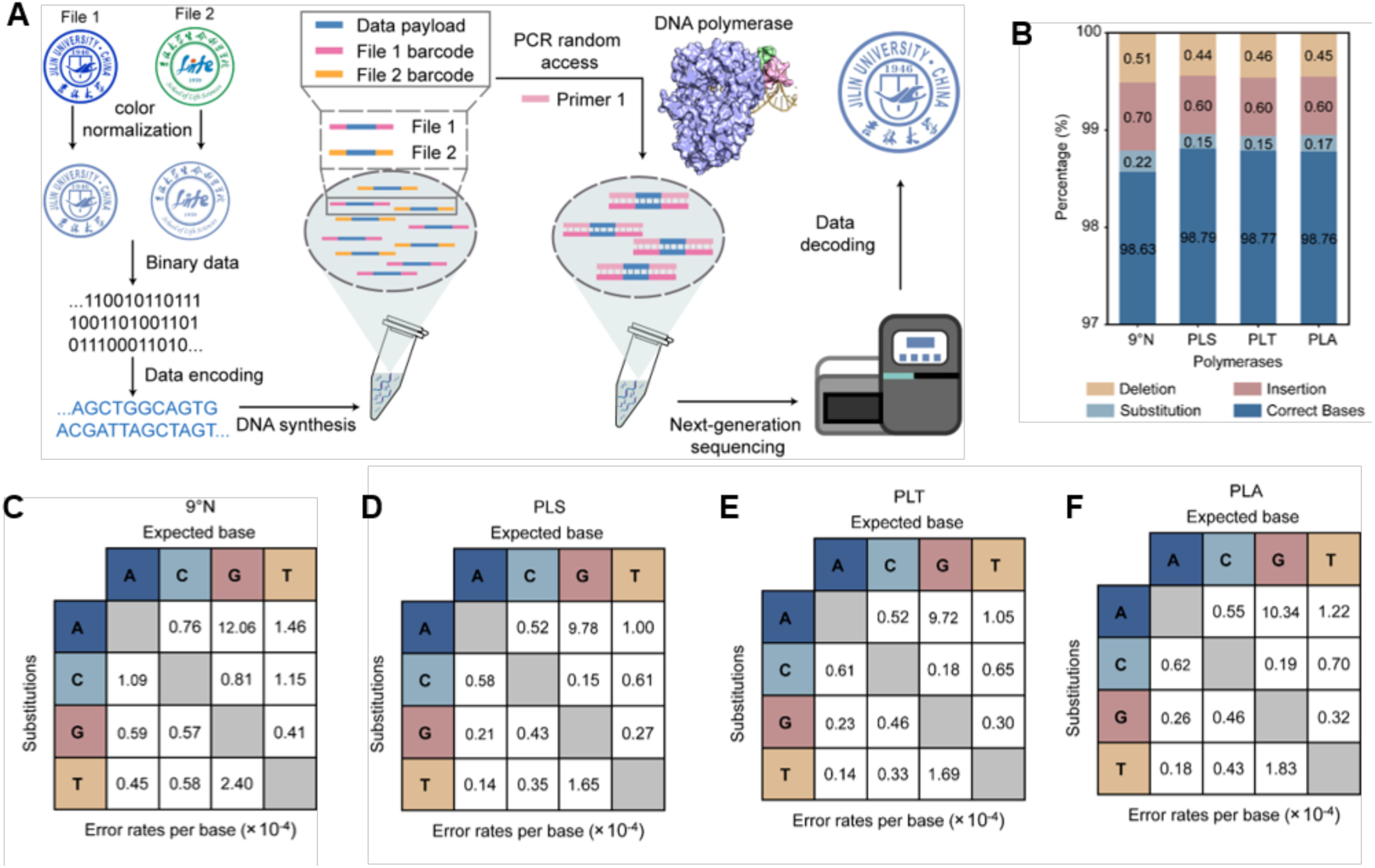
Information retrieval using DNA polymerases. (**A**) Schematic representation of data retrieval using orthogonal barcode. (**B**) Analysis of error types and frequencies in DNA polymerases amplification. (**C-F**) Base substitution matrices for 9°N (**C**), PLS (**D**), PLT (**E**), and PLA (**F**) DNA polymerases.

### 3.4 Chimeric DNA polymerase enabled modular DNA movable type storage

Inspired by ancient Chinese movable type printing, DNA movable type storage offers significant advantages in cost reduction and operational flexibility [**13–16**]. This strategy assembles reusable, prefabricated DNA elements to encode information, thereby minimizing the requirement for repeated *de novo* DNA synthesis and substantially reducing both costs and labor while enabling modular information storage. In this system, short DNA fragments, termed DNA movable types (DMTs) were enzymatically assembled into DNA movable type blocks (DMTBs) using chimeric DNA polymerases. Each DMTB served as an encoding unit for a Chinese character composed of three parts: an address movable type (AMT) for sequence positioning, a checksum movable type (CMT) for error correction, and a payload movable type (PMT) for data encoding. The AMTs employed a three-level addressing system (A1∼4, B1∼4, C1∼4), enabling precise positioning of individual data segments and supporting random file access. This design also improved synthesis efficiency. For example, storing 1,000 DMTBs with a single-level address would require 1,000 distinct address sequences. In contrast, three-level addressing system could achieve the same capacity through combinatorial assembly, where each of the three address levels (A, B and C) contained 10 distinct modules. This hierarchical design allowed 10 × 10 × 10 = 1,000 unique address combinations using only 30 synthesized sequences in total. To allow *in vivo* storage, the assembled DMTBs were ligated into the pUC19 plasmid at BamHI and HindIII sites and transformed into *E. coli*. A classical Chinese poem “wang lu shan pu bu” (Pinyin, meaning Gazing at the Waterfall on Mount Lu), composed by the poet Li Bai during Tang Dynasty, was selected for encoding. Each Chinese character, punctuation marks and carriage return in the poem was treated as an independent movable type and encoded into a distinct DMTB, where DMTB1 corresponded to the character “wang”, DMTB2 to “lu”, DMTB3 to “shan” and so on. Each DMTB of the poem was 145 nt in length, comprising a 60 nt AMT, a 20 nt CMT, and a 50 nt PMT, connected by 5 nt linker sequences. This approach conceptually followed by the classical Chinese movable type printing system, where each character served as a reusable and position-specific type block. To efficiently assemble a complete DMTB from its constituent parts (AMT, CMT and PMT), we performed one-pot overlap PCR using PLS DNA polymerase, which was selected for its superior performance. Increasing the concentration of PLS DNA polymerase significantly enhanced the assembly efficiency. At an optimal concentration of 8.0 pM, the AMT, CMT and PMT were successfully assembled into a full-length DMTB. To enable efficient integration into the pUC19 plasmid, homologous sequences corresponding to regions flanking the BamHI and HindIII restriction sites were appended to both ends of each DMTB, thereby facilitating homologous recombination. Accordingly, all 44 DMTBs were subsequently synthesized using identical conditions and inserted into linearized pUC19 plasmids (**Fig. S11**). These resulting DMTB plasmids were transformed into *E. coli*, establishing a plasmid-based DMTB library that supported long-term data preservation within living cells. To reconstruct the original file from the *in vivo* stored DNA, all plasmid samples were sequenced, and the resulting reads were aligned to their corresponding reference sequences for decoding. The high fidelity of the PLS DNA polymerase used in the enzymatic assembly ensured that each DMTB retained its designed information, allowing accurate recovery of address, error correction, and data payload during assembly. The successful sequencing and decoding of all DMTBs enabled complete reassembly of the poem with perfect data integrity, demonstrating that PLS DNA polymerases played a key role in preserving sequence accuracy during both enzymatic assembly and plasmid-based storage. To further investigate the fidelity of *in vivo* storage, we analyzed error types of all 44 DMTBs, including insertions, deletions and substitutions. By decoding the sequencing data of all DMTBs, the original file was successfully reconstructed. During file reconstruction, errors such as deletions, substitutions, and insertions were observed in the sequencing results of DMTBs assembly. As shown in **Table S6**, over 99.9% of the reads were perfectly decoded, with only approximately 0.092% of the reads containing errors. Among the observed errors, deletions were the most frequent (0.0764%), followed by substitutions (0.0117%) and insertions (0.0039%). Deletion errors primarily occurred in the AMT and CMT regions, substitution errors were mainly located in the AMT, while insertion errors were concentrated in the CMT. This distribution of errors was likely caused by the sequencing orientation, which proceeded from AMT to PMT. As sequencing errors were more common at the initial positions of reads, this might explain the higher error rates observed in the AMT and CMT regions. Despite these low-frequency errors, the embedded CMT enabled accurate error correction, ensuring complete and accurate reconstruction of the original file.

To validate the reusability of DNA movable types, existing DMTBs plasmids stored in *E. coli*. were used as templates for overlap PCR to reconstruct new DMTBs (ReDMTBs) without *de novo* DNA synthesis. Specifically, five ReDMTBs were constructed by recombining the AMT of one DMTB with the CMT and PMT of another. For example, ReDMTB1 was generated by combining the AMT of DMTB1 (encoding the character “wang”) with the CMT and PMT of DMTB4 (encoding the character “pu”). This recombinant DMTB preserved the original positional information of “wang” while encoding the character “pu”. Similarly, ReDMTB2 was formed from the AMT of DMTB2 (“lu”) and the CMT and PMT of DMTB5 (“bu”), resulting in ReDMTB2 encoding “bu” with the address of “lu”. Following this approach, ReDMTB3 combined the AMT of DMTB3 (“shan”) with the CMT and PMT of DMTB38 (“si”), ReDMTB4 used the AMT of DMTB4 (“pu”) and the CMT and PMT of DMTB39 (“yin”), and ReDMTB5 was constructed from the AMT of DMTB5 (“bu”) and the CMT and PMT of DMTB40 (“he”). Through this modular recombination strategy, the original phrase “wang lu shan pu bu” (Gazing at the Waterfall on Mount Lu) to a new phrase “pu bu si yin he” (The Waterfall Looks Like the Milky Way), while preserving positional information through the original AMTs. Sequencing results confirmed that each ReDMTB retained the positioning address of one parent DMTB and the information content of the other (**Fig. S12**-**S16**. These findings demonstrated the feasibility of using chimeric DNA polymerase-based overlap PCR to recombine prefabricated DNA blocks, enabling low-cost, flexible rewriting of stored data without *de novo* synthesis. Moreover, this approach highlighted the potential of DNA movable type storage for dynamic content reconfiguration through the selective reuse of modular encoding units.

To expand the capability of DNA movable type storage, we further explored the sequential and multi-fragment assembly of prefabricated DNA blocks for constructing longer and more complex information sequences. Traditional oligonucleotide-based storage systems are often constrained by synthesis length limits and fragmented sequence pools, making it difficult to assemble extended data structures. In contrast, DNA movable types can be programmatically ordered and enzymatically assembled to form long and continuous strands. To demonstrate this advantage, PMT4 (encoding “pu”), PMT 5 (“bu”), PMT 38 (“si”), PMT 39 (“yin”), and PMT 40 (“he”) were selected and sequentially assembled using polymerase cycling assembly (PCA) (**Fig. S17**). Sequencing results validated the successful assembly and accurate decoding of the five PMTs into the intended phrase “pu bu si yin he” (meaning “The Waterfall Looks Like the Milky Way”) (**Fig. S18** and **S19**). However, PCA requires stringent control of melting temperatures, sequence composition, and secondary structures, limiting its scalability for large DNA movable type libraries. To overcome these limitations, we employed a multi-enzyme assembly strategy that integrated Golden Gate and Gibson assembly methods. Golden Gate assembly enabled precise, seamless ligation of multiple DNA fragments *via* type IIS restriction sites, while Gibson assembly facilitated the insertion of assembled fragments into plasmid backbones through homologous recombination. In this approach, DMTBs were first amplified from existing information plasmid using PLS DNA polymerase. During amplification, BsaI recognition sites were introduced at the termini of each DMTB. BsaI enzymes cleaved beyond its recognition sequence to generate specific sticky ends while eliminating the recognition sites, thereby enabling seamless cloning and directional ligation for efficient assembly. Subsequently, T4 DNA ligase was employed to sequentially ligate the sticky ends of multiple DMTBs, producing long DNA fragments. These fragments were designed with homologous regions flanking both ends to facilitate recombination into linearized vectors. The assembled constructs were then transformed into *E. coli* for sequencing and long-term storage. Using this strategy, we successfully assembled nine DMTBs into a single DNA movable type unit (DMTU) plasmid. The order of DMTBs assembly was defined by the unique restriction patterns introduced by PLS DNA polymerase, enabling accurate connection of adjacent storage blocks. Ultimately, all 44 DMTBs representing the full poem were grouped into five DMTUs. Specifically, DMTU1 comprised DMTB1 to DMTB9, DMTU2 included DMTB10 to DMTB18, DMTU3 was assembled from DMTB19 to DMTB27, DMTU4 from DMTB28 to DMTB36, and DMTU5 from DMTB37 to DMTB44. Agarose gel electrophoresis verified the successful assembly of the expected DNA fragments. (**Fig. S20**). Furthermore, these five DMTUs were sequentially assembled into a complete DNA fragment (∼4.4 kb), referred to as the DMTF (DMT-File), which encoded the full content of the classical Chinese poem (**Fig. S21**). Sequencing analysis confirmed complete data recovery, validating the reliability of the DNA movable type strategy. This enzymatic approach enabled the scalable and accurate assembly of long DNA sequences from modular, reusable units. Compared to traditional chemical synthesis, which is generally limited to sequences under 200 nt, this DNA polymerase-based assembly method allowed the production of kilobase-scale DNA strands, significantly enhancing data capacity per molecule while reducing synthesis costs.

### 3.5 Robustness of DNA Movable Type Storage *in vivo*

Long DNA fragments in engineered bacteria are inherently prone to loss, translocation, and spontaneous mutations during extended passage. To evaluate the long-term stability of the *in vivo* DNA movable type storage system, six randomly selected DMTBs and DMTUs were cultured for 100 generations (**Fig. 5A**). All retrieved sequences remained identical to the reference, confirming that information stored in *E. coli* remained stable throughout serial passaging (**Fig. 5B**). Synthetic oligonucleotide pools degrade over time and must be resynthesized. In contrast, *in vivo* storage leverages cell replication to maintain data integrity without additional synthesis. This intrinsic capability enables autonomous restoration of data lost due to degradation or frequent retrieval. To assess robustness under repeated retrieval, DMTB5 and DMTU2 were continuously cultured over a 72-h sampling experiment (**Fig. 5C**). At 12 h-intervals, fixed volumes of bacterial culture (25 mL, 12.5 mL, or 5 mL from a total volume of 50 mL) were extracted for sequencing, and fresh medium was replenished to maintain growth. After each sampling, bacterial density temporarily declined but consistently recovered within 12 h, indicating stable cell proliferation and sustained maintenance of the stored DNA (**Fig. 5D**). Moreover, sequencing results across all time points showed error-free data retrieval, indicating that in vivo storage stably preserved digital information even under frequent access. Collectively, these results demonstrated that the polymerase-assembled, plasmid-based DNA movable type system achieved stable, high-fidelity data storage in living cells, offering a biologically regenerative and cost-efficient alternative to conventional chemical synthesis-based DNA storage platforms.

**Fig. 5.**
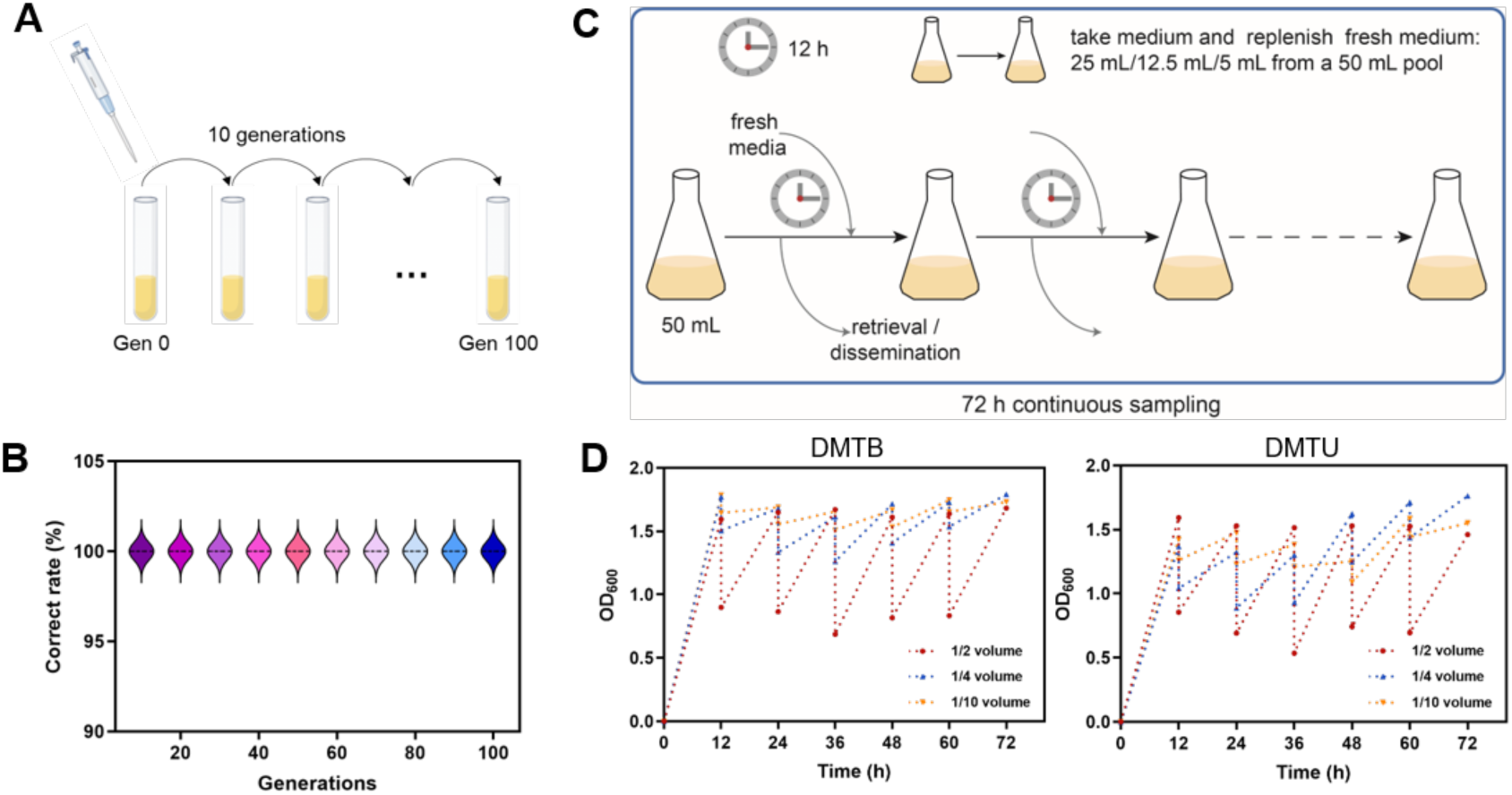
Storage stability of long-term DNA movable type storage *in vivo*. (**A**) Bacterial passaging experiment. (**B**) Sequencing analysis of DMTB- and DMTU-engineered bacteria after serial passaging. (**C**) Sampling scheme for continuous bacterial growth. (**D**) OD600 measurement of DMTB- and DMTU-engineered bacteria under different sampling volumes.

## 4. Discussion and conclusion

DNA polymerases are essential enzymes in DNA-based information storage systems, enabling high-fidelity amplification, precise file retrieval, modular sequence assembly, and supporting long-term preservation through accurate replication [**39**]. However, most current storage architectures, particularly those relying on synthetic oligonucleotide pools, primarily emphasize chemical synthesis while insufficiently leveraging the catalytic functions of DNA polymerases. This reliance on *de novo* chemical synthesis leads to short sequence lengths, high synthesis costs, and limited flexibility in data reconfiguration, which in turn constrains the scalable and practical application of DNA storage. To address these challenges, we previously demonstrated the feasibility of encoding diverse data types, including images, text, and videos, into DNA oligonucleotide pools using the thermostable 9°N DNA polymerase [**20**]. Compared with widely used commercial polymerases, 9°N exhibited higher fidelity, which effectively reduced base substitution errors during the synthesis process. Despite its high fidelity, 9°N lacked strong processivity due to the absence of accessory proteins, restricting its capacity for long-fragment assembly. Thus, these limitations motivated us to develop an engineered polymerase-driven DNA movable type storage strategy that utilized DNA polymerases to assemble reusable prefabricated DNA modules. To improve enzymatic assembly efficiency, we enhanced DNA polymerase processivity by fusing the non-replicative 9°N DNA polymerase with archaeal DNA binding proteins (Sso7d, Sac7d, Sto7d) *via* flexible peptide linkers to preserve structural integrity. Compared with wild-type 9°N DNA polymerase, these chimeric polymerases exhibited significantly improved processivity, thermal stability, salt tolerance, fidelity and the ability to amplify long DNA fragments. Molecular dynamics simulations revealed that fused DNA binding proteins increased polymerase-template interactions *via* minor groove contacts, mimicking the function of sliding clamps in maintaining binding stability. Notably, differences in binding strength among the chimeric enzymes led to distinct functions. Although PLA exhibited the strongest DNA binding affinity among the chimeric polymerases, it showed reduced catalytic efficiency, likely due to excessive stabilization of the enzyme/template complex. This strong binding might hinder the DNA strand translocation to DNA polymerase, thereby limiting the polymerase’s ability to efficiently incorporate successive nucleotides during DNA synthesis. In contrast, PLS DNA polymerase demonstrated a balanced DNA binding affinity that effectively stabilized the complex without restricting translocation, thereby achieving both high fidelity and long-fragment amplification capability. In addition to improving DNA synthesis performance, the engineered polymerases also enhanced random access fidelity during data retrieval. We designed orthogonal barcode primers with high sequence dissimilarity and minimal secondary structure to enable multiplexed access to distinct data files stored in a shared pool. Using this strategy, chimeric polymerases accurately retrieved image files representing the “Jilin University” and “School of Life Sciences” logos from a mixed oligonucleotide pool. Moreover, compared with wild-type 9°N, PLS DNA polymerase exhibited significantly reduced substitution error rates and increased correct read proportion, demonstrating its enhanced fidelity and primer specificity in random-access retrieval from mixed DNA pools. This improved retrieval accuracy demonstrated the pivotal role of engineered DNA polymerases in facilitating sequence-specific file access in DNA-based information storage system.

Motivated by the enhanced catalytic properties of the engineered DNA polymerases in both amplification and addressing, we next applied them to construct a DNA movable type storage system that enabled enzymatic modular assembly and flexible data rewriting. In this system, prefabricated address, checksum, and payload modules were enzymatically assembled into DMTBs *via* polymerase-mediated overlap extension PCR. Compared with traditional oligonucleotide-based approaches, this design enabled accurate information writing without requiring chemical ligation or extensive repeated synthesis of oligonucleotides, thereby reducing cost and enhancing flexibility for DNA data construction. To demonstrate the modular rewriting capacity of this system, we recombined existing DMTBs into recombinant units (ReDMTBs) by selectively replacing checksum and payload modules while preserving their original address sequences. This modular recombination strategy enabled content rewriting at the sequence level by flexibly reassembling address, checksum, and payload units, allowing efficient reuse of prefabricated blocks without additional synthesis. Furthermore, by establishing an *in vivo* library of DMTBs, we demonstrated that new information could be rapidly constructed through selective combination of existing components, highlighting the system’s scalability and potential for dynamic, low-cost data rewriting. To support the construction of longer storage units, DMTBs were further assembled into multi-block DNA movable type units (DMTUs). Initially, PCA was employed to combine multiple payload modules into sentence-level structures. Using this method, we successfully assembled five PMTs into a continuous DNA fragment encoding the new phrase “The Waterfall Looks Like the Milky Way”, demonstrating the potential of modular assembly for structured data synthesis. However, PCA was limited by stringent sequence requirements and thermal constraints, which restricted its scalability for assembling more complex or higher-diversity sequence libraries [**40**, **41**]. To overcome these limitations, we utilized a hybrid enzymatic assembly strategy that combined Golden Gate assembly with Gibson assembly. Golden Gate assembly enabled scarless and directional joining of multiple fragments, while Gibson assembly offered greater flexibility for integrating larger or structurally diverse constructs into plasmids [**42**, **43**]. Notably, PLS DNA polymerase was used to amplify full-length DMTBs directly from plasmids containing the stored information. During this amplification, primers were designed to introduce either Type IIS restriction sites or homologous overlaps at the termini of each fragment, enabling compatibility with both assembly methods. Meanwhile, the high fidelity of PLS polymerase ensured accurate sequence replication, minimizing errors during the multi-fragment construction process. Using this strategy, nine DMTBs were first assembled into a single plasmid to form one DNA movable type unit (DMTU), validating the feasibility of multi-block enzymatic assembly. Subsequently, all 44 DMTBs encoding an entire Chinese poem were grouped into five DMTUs. These DMTUs were further assembled into a full-length file (DMTF) of ∼4.4 kb and stored in engineered plasmids. This kilobase-scale, sequence-defined DNA construct demonstrated the potential of enzymatic assembly in overcoming the length limitations of chemical synthesis. Sequencing verification of all assembled DMTUs and the final DMTF confirmed complete sequence fidelity, highlighting the crucial role of PLS DNA polymerase in ensuring accuracy during multi-fragment amplification and integration.

To evaluate the long-term stability and accessibility of this storage system, we introduced the assembled DMTUs into *E. coli* and assessed data retention over 100 generations and repeated sampling. Sequencing results confirmed that all stored information remained intact and error-free, exhibiting DNA polymerase-assembled DNA movable types could be faithfully maintained through bacterial replication. These findings indicated that enzymatically-assembled DNA blocks could serve as stable and regenerable storage units and enabled the construction of sustainable DNA movable type libraries in living cells for long-term data preservation.

Collectively, our findings demonstrated the central role of engineered DNA polymerases as versatile molecular tools for DNA-based information storage. Their enhanced processivity and fidelity were essential not only for constructing modular, long-fragment DNA movable type systems, but also for achieving accurate file retrieval from mixed oligonucleotide pools. Moreover, the engineered polymerase-driven system reduced synthesis burden, enabled enzymatic rewriting, and ensured long-term stability in living hosts, offering a robust foundation for scalable and sustainable DNA data storage. In the future, the integration of multiple enzymes into microfluidic platforms might enable automated, cascade DNA assembly and rewriting workflows, further improving efficiency and throughput of enzymatic DNA storage systems.

## 5. Acknowledgements

The work was supported by the National Key R&D Program of China (2020YFA0907000), the National Natural Science Foundation of China (U24A20365, 32271319, 32201030 and 32471315), the Science and Technology Department of Jilin Province (20240402035GH), the Development and Reform Commission of Jilin Province (2023C015 and 2024C013-8) and the Fundamental Research Funds of the Central Universities, China (2024-JCXK-11).

## 7. Data availability

Some data originally included in this study are not displayed in the current version of the manuscript in accordance with bioRxiv policies and are available from the corresponding author upon reasonable request.

