## Supplementary information for "Engineering chimeric DNA polymerases for DNA movable type storage"

*Corresponding authors.

**Molecular dynamics (MD) simulation**

The 9°N DNA polymerase was modeled using PDB 4K8X for the protein structure and PDB 5OMV for divalent cation coordination. The Sso7d and Sac7d DNA binding proteins were modeled using PDB 1BNZ and PDB 1AZQ, respectively. The Sto7d DNA binding protein was modeled using the AlphaFoldDB Q96X56. Gaussian accelerated molecular dynamics (GaMD) simulation is an enhanced sampling technique that involves adding a harmonic boost potential to smoothen the system's potential energy surface. To enhance conformational sampling, GaMD simulations of four model systems of 9°N DNA polymerases and three chimerical DNA polymerases were performed using the graphic processing unit (GPU) version of Amber 20 [1]. The DNA was described using the Amber OL15 force field. $K^+$ and $Cl^-$ were added to the protein surface to neutralize the total charges of the systems. The resulting systems were solvated in a rectangular box of TIP3P10 waters extending up to a minimum cutoff of 10 Å from the protein boundary. The Amber ff14SB force field was employed for the protein in all the MD simulations [2]. The system energy minimization with the steepest descent algorithm and conjugate gradient algorithm was executed to eliminate atomic collisions in the initial structure (5000 steps) [3]. Then, the three systems were gradually heated from 0 K to 345 K under an NVT ensemble, and then relaxed under an NPT ensemble. Minimization, heating, and equilibration were completed, and production simulations were performed for 500 ns. The trajectories from the simulations of the binary and ternary structures were analyzed using CPPTRAJ [4] and the VMD [5] program for calculating root-mean-square deviation

(RMSD), root-mean-square fluctuation (RMSF), Radius of gyration (Rg), and contact fraction analysis. All molecular graphics showing structures were prepared with PyMOL (https://pymol.org/pymol/).

**Binding energy calculation**

The Poisson-Boltzmann or generalized Born and surface area continuum solvation (MM/PBSA and MM/GBSA) methods, which has been widely used to estimate the free energy of the binding of ligand to biological macromolecules. In this study, MM/PBSA and MM/GBSA was employed to calculate the binding energy of different chimera DNA polymerase-DNA template complexes. The free energy of each molecule was calculated as follows: $\Delta G_{binding}=G_{complex}-(G_{protein}+G_{ligand})$. Here, $G_{complex}$, $G_{protein}$, and $G_{ligand}$ are the free energy of the DNA polymerase-DNA complex, the free energy of the DNA polymerase, and the free energy of DNA, respectively. All binding energies were calculated with the g_mmpbsa tool.

**Orthogonal DNA primer design**

To enable specific DNA barcode, the primer optimization algorithm was performed to design orthogonal primer pairs. We chose two pairs of orthogonal DNA primers from primer library designed above [6]. The principles for designing a primer library were as follows. The process started with a randomly generated 20-mer primer, which was scored against several design criteria: (1) no long homopolymer regions ($\leqslant$ three consecutive A/T, $\leqslant$ two consecutive G/C); (2) $\leqslant$ 4 bases of self-complementarity and $\leqslant$ 10 bases of inter-sequence complementarity; (3) 45%-55% GC content; and (4) a minimum Hamming distance of 6 from all other primers. If a sequence violated a design criterion, +1 penalty was assigned to the corresponding base, and the total primer score was computed by summing all base-level scores. Primers with lower total scores were selected. Further filtering was performed based on secondary structure and melting temperature. Candidates were then screened using BLAST to ensure no inter-primer similarity >12 bp. Additionally, a “collision check” aligned data payload against the primers to prevent any >12 bp overlaps, ensuring sequence orthogonality. Using this approach, JLU-F/R and SLS-F/R primer pairs were designed to retrieve data corresponding to the “Jilin University” and “School of Life Sciences” logos, respectively.

**Data sequencing and analysis**

Sequencing and data analysis were as previously described [7]. Raw sequencing data underwent initial quality control using Fastp [8], followed by adapter trimming with default parameters in Trimmomatic [9]. Overlapping reads were assembled into single contigs *via* barcode-guided merging using SeqPrep (https://github.com/jstjohn/SeqPrep) to determine oligonucleotide positions. These contigs were then aligned to the reference index employing BWA-MEM [10]. Post-alignment error rates were calculated based on variant calling using the samtools [11].

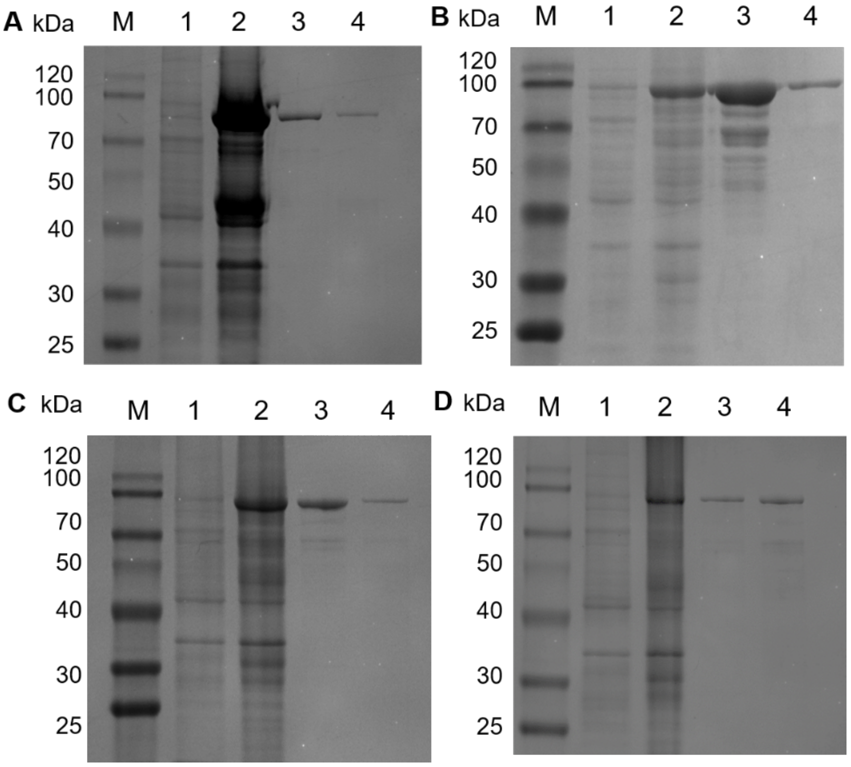


**Fig. S1.** SDS-PAGE analysis of DNA polymerases expressed in *E. coli* BL21 (DE3). (**A**) 9°N, (**B**) PLS, (**C**) PLT, and (**D**) PLA. M: Marker; Lane 1: Cell lysates without IPTG induction; Lane 2: Cell lysates with IPTG induction; Lane 3: supernatants after Ni-affinity purification; Lane 4: supernatants after HisTrap Q column purification.

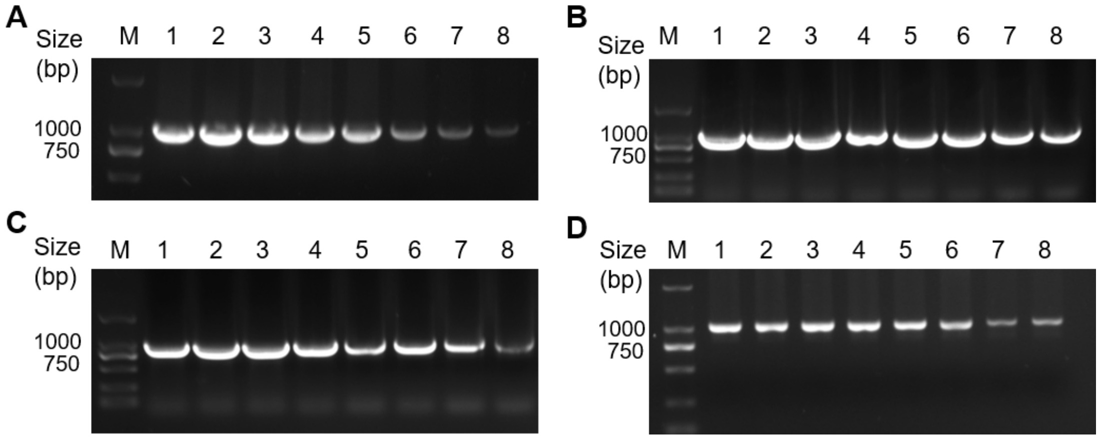


**Fig. S2.** Thermal stability analysis of DNA polymerases. (**A**) 9°N, (**B**) PLS, (**C**) PLT, and (**D**) PLA. M: Marker; Lane 1-8: enzymes incubated at 95 °C for 0, 5, 10, 20, 40, 60, 90, and 120 min, respectively.

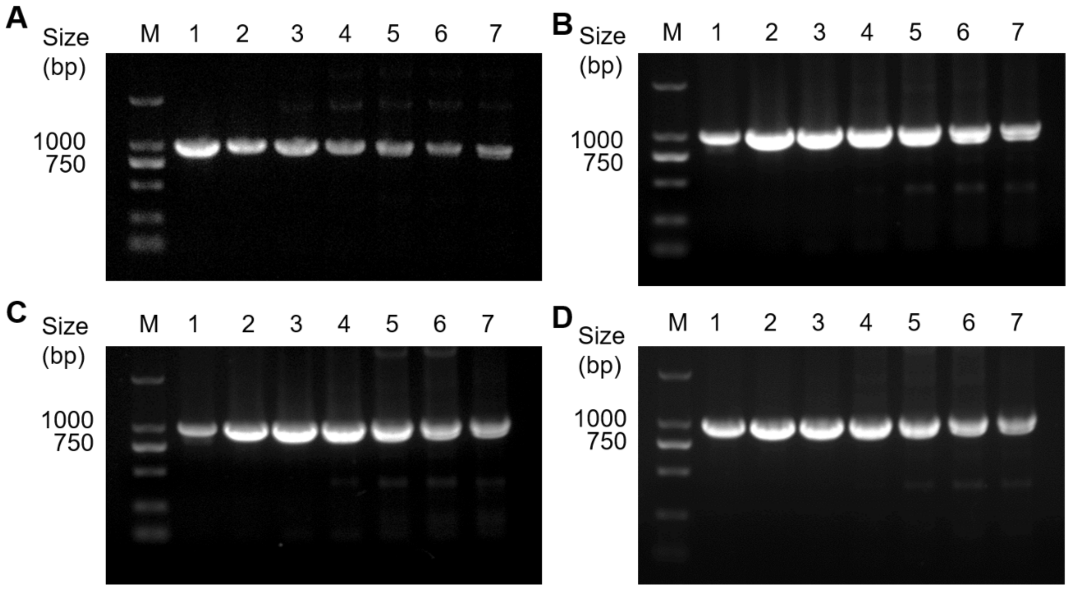


**Fig. S3.** Salt tolerance analysis of DNA polymerases. (**A**) 9°N, (**B**) PLS, (**C**) PLT, and (**D**) PLA. M: Marker; Lane 1-7: enzymes incubated at KCl concentration of 0, 10, 20, 30, 40, 50, and 60 mM, respectively.

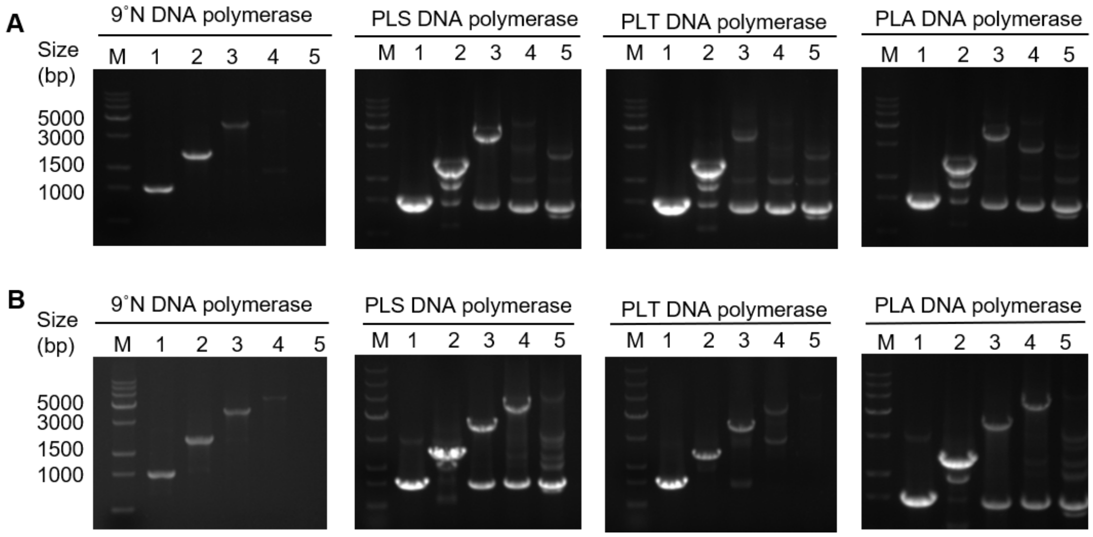


**Fig. S4.** Evaluation of PCR efficiency using λDNA as template under different thermocycling conditions in the absence of KCl. (**A**) PCR with 30 s extension at 72 °C; (**B**) PCR with 60 s extension at 72 °C. M: DNA marker; Lanes 1-5: target amplicons of 1, 2, 4, 6, and 8 kb, respectively.

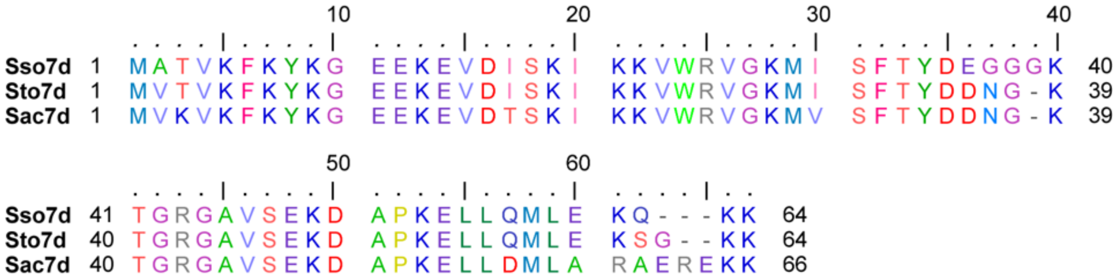


**Fig. S5.** Multiple sequence alignment of DNA binding proteins Sso7d, Sto7d, and Sac7d.

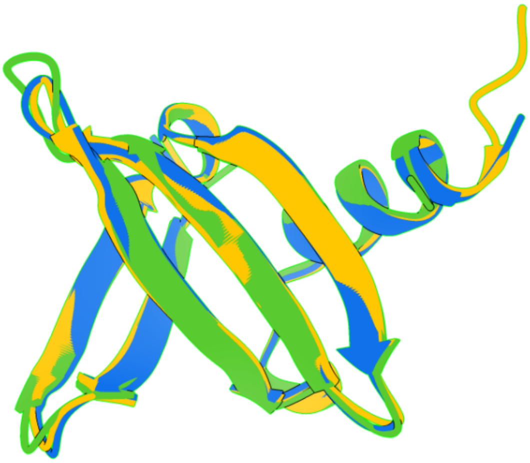

**Fig. S6.** Structural alignment of DNA binding proteins Sso7d (green), Sto7d (blue), and Sac7d (orange) to compare their overall conformational similarities.

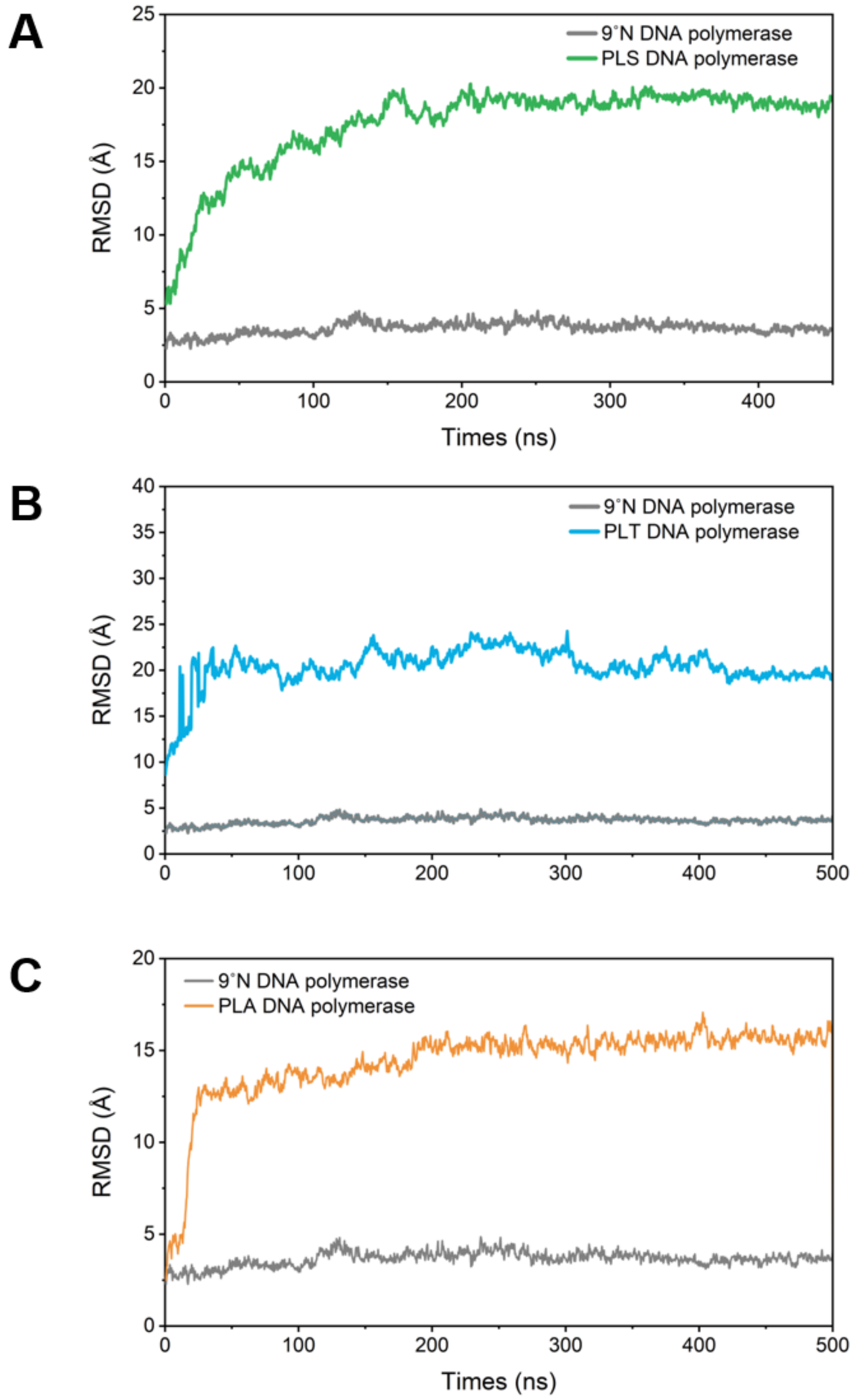


**Fig. S7.** Root-mean-square deviation (RMSD) of the polymerase backbone in chimeric DNA polymerase/dsDNA complexes.

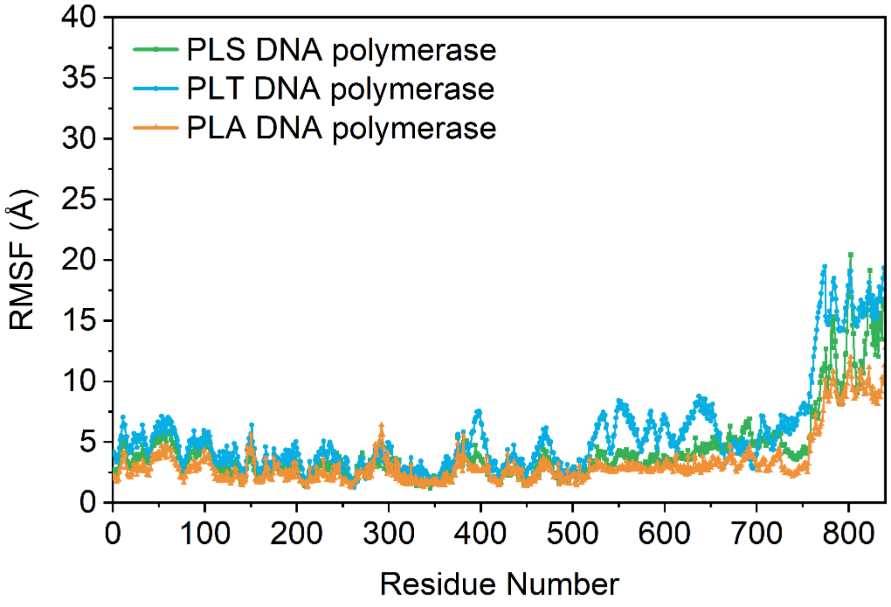


**Fig. S8.** Root-mean-square fluctuation (RMSF) of residues in chimeric DNA polymerase systems during molecular dynamics simulations.

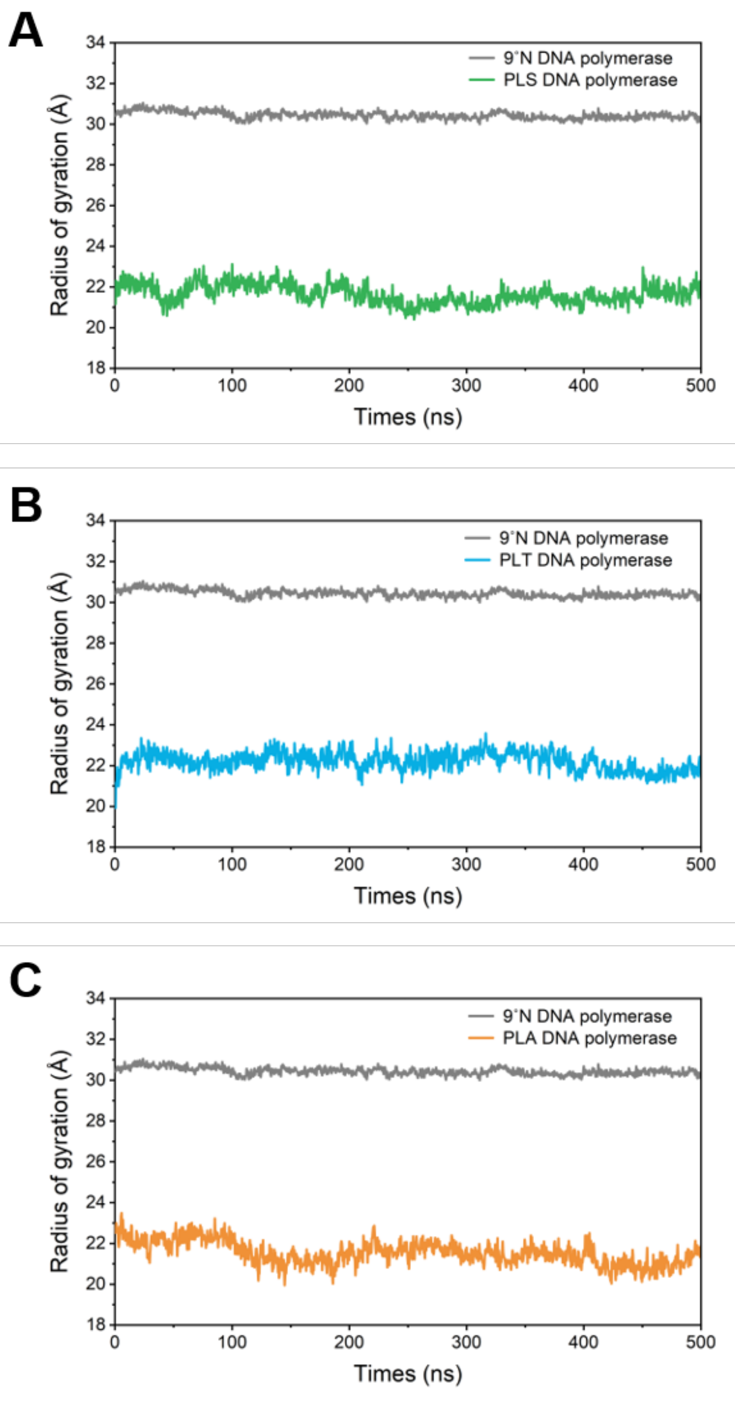


**Fig. S9.** Radius of gyration (Rg) profiles of chimeric DNA polymerases with fused DNA binding proteins during molecular dynamics simulations.

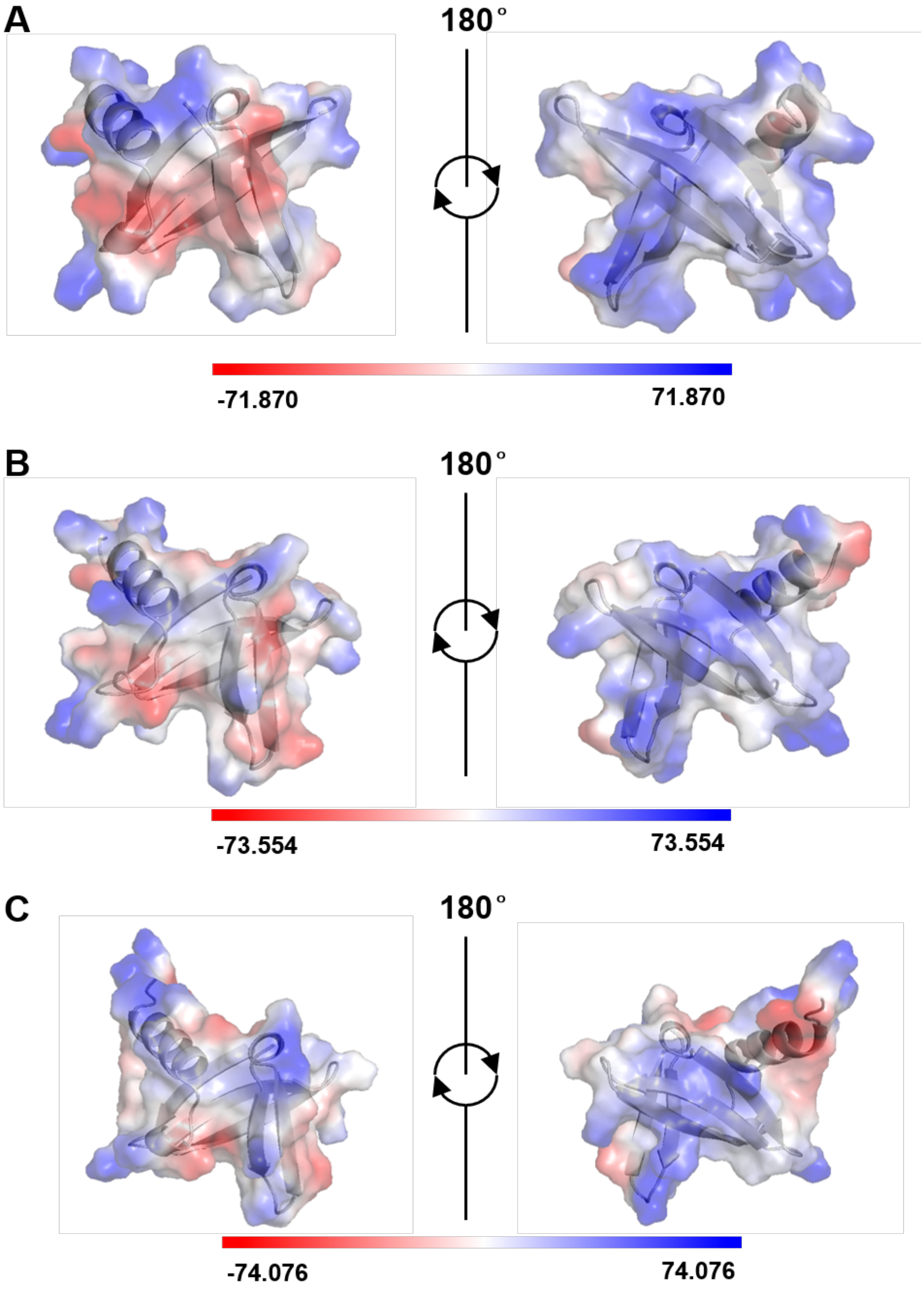


**Fig. S10.** Surface electrostatic potential of the DNA binding protein in (**A**) PLS, (**B**) PLT, and (**C**) PLA DNA polymerases. The right panels showed the regions of the DNA binding proteins that contacted dsDNA. Electrostatic potential was colored as positive (blue), negative (red), and neutral (white).

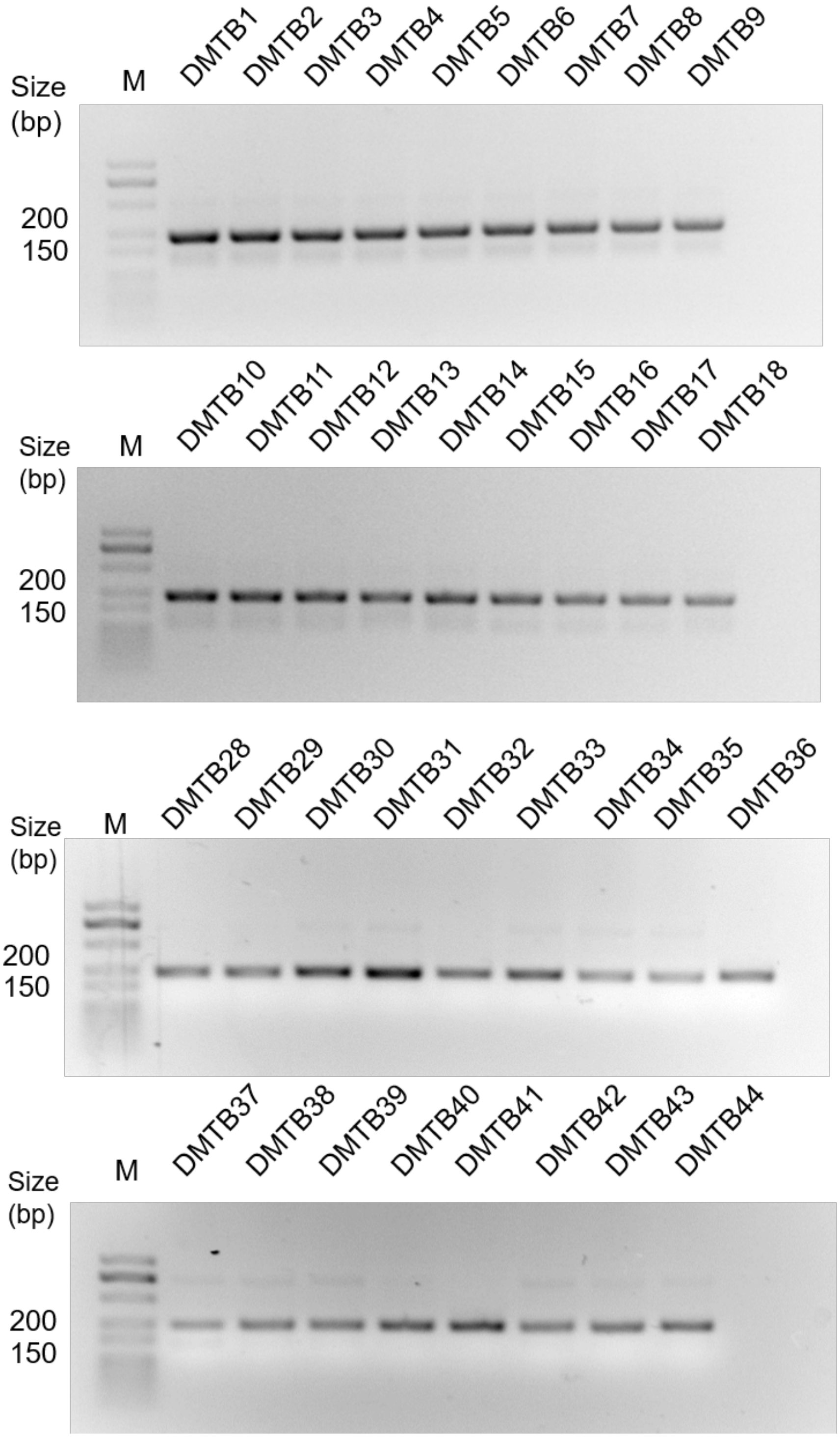


**Fig. S11.** Gel electrophoresis analysis of 44 DMTBs assembled using PLS DNA polymerase. M: Marker.

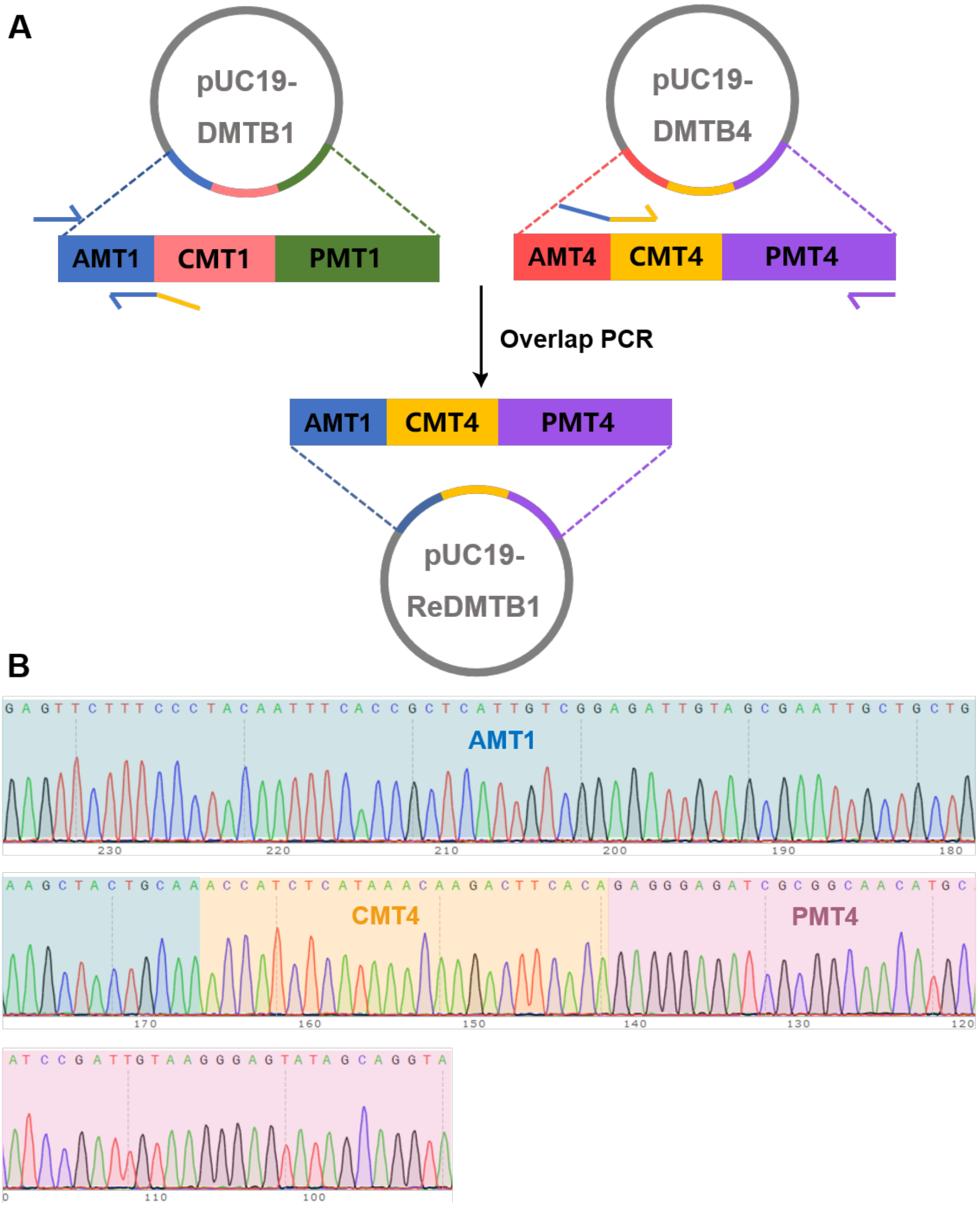


**Fig. S12.** Construction of the ReDMTB1. (A) ReDMTB1 was constructed by combining the address module (AMT) of DMTB1, which encoded the character "wang", with the checksum and payload modules (CMT and PMT) of DMTB4, which encoded the character "pu". This recombinant block retained the positional identity of "wang" while representing "pu". (**B**) Sequencing analysis of ReDMTB1.

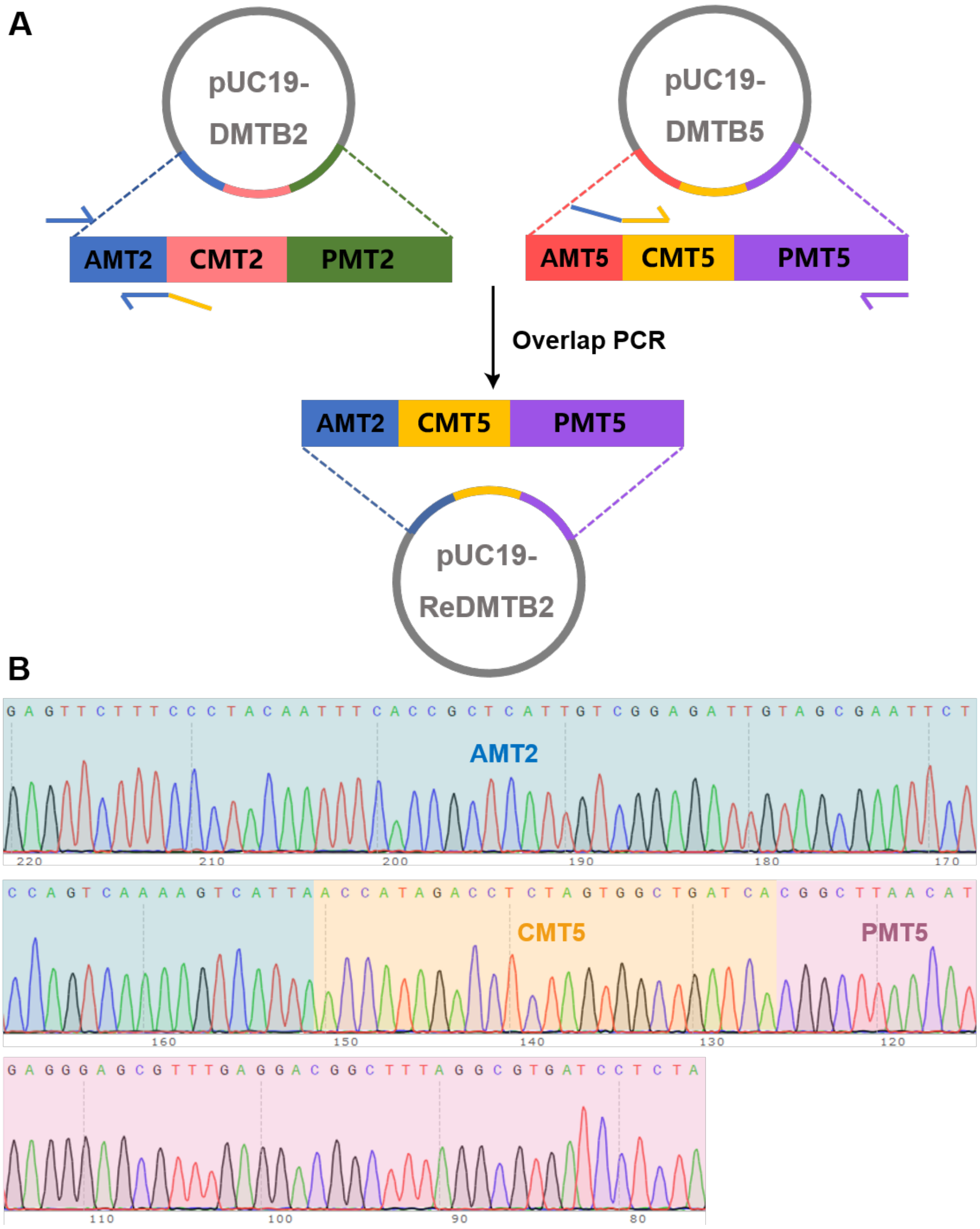


**Fig. S13.** Construction of ReDMTB2. (**A**) ReDMTB2 was constructed by combining the address module (AMT) of DMTB2, which encoded the character "lu", with the checksum and payload modules (CMT and PMT) of DMTB5, which encoded the character "bu". This recombinant block retained the positional identity of "lu" while representing "bu". (**B**) Sequencing analysis of ReDMTB2.

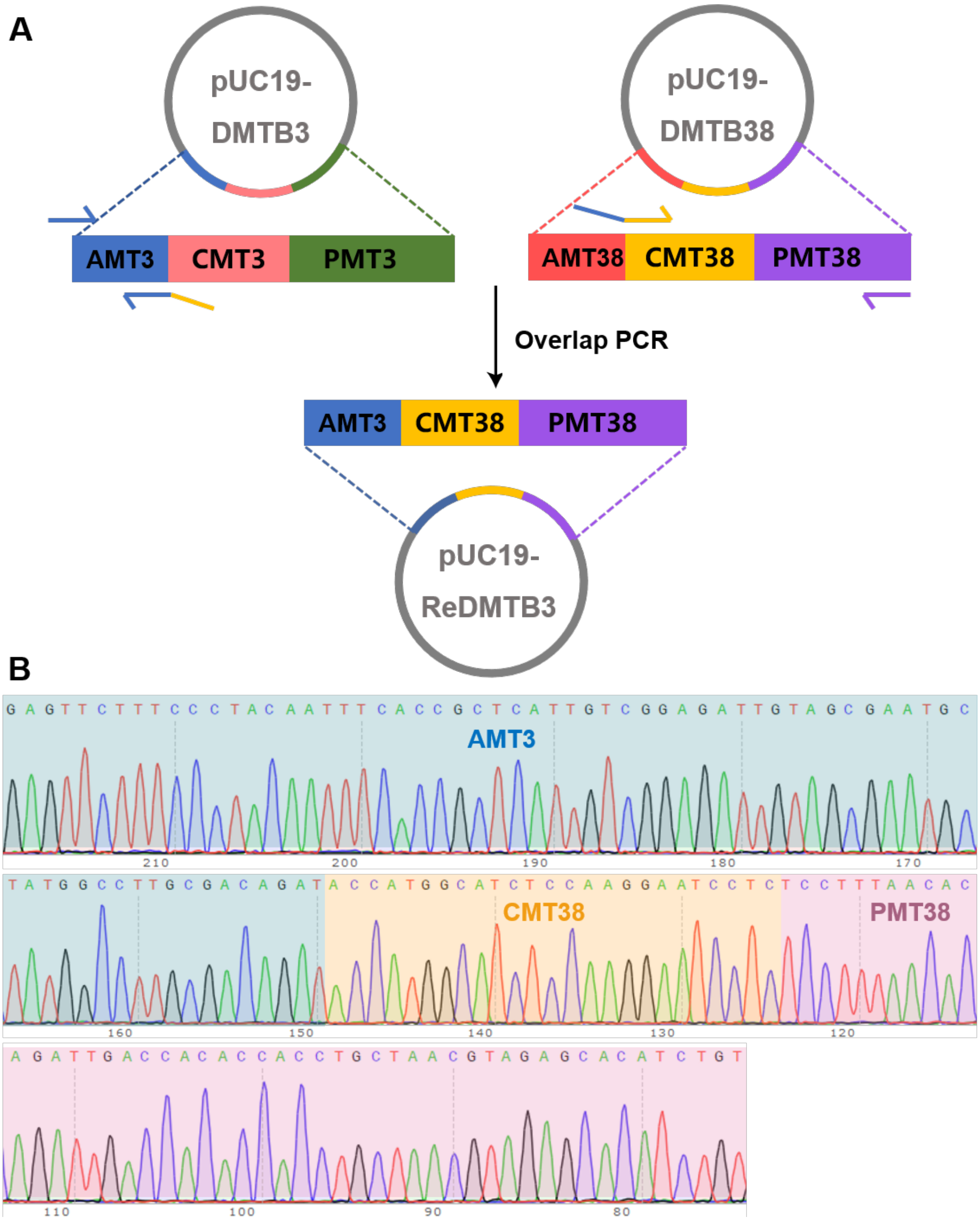


**Fig. S14.** Construction of ReDMTB3. (**A**) ReDMTB3 was constructed by combining the address module (AMT) of DMTB3, which encoded the character "shan", with the checksum and payload modules (CMT and PMT) of DMTB38, which encoded the character "si". This recombinant block retained the positional identity of "shan" while representing "si". (**B**) Sequencing analysis of ReDMTB3.

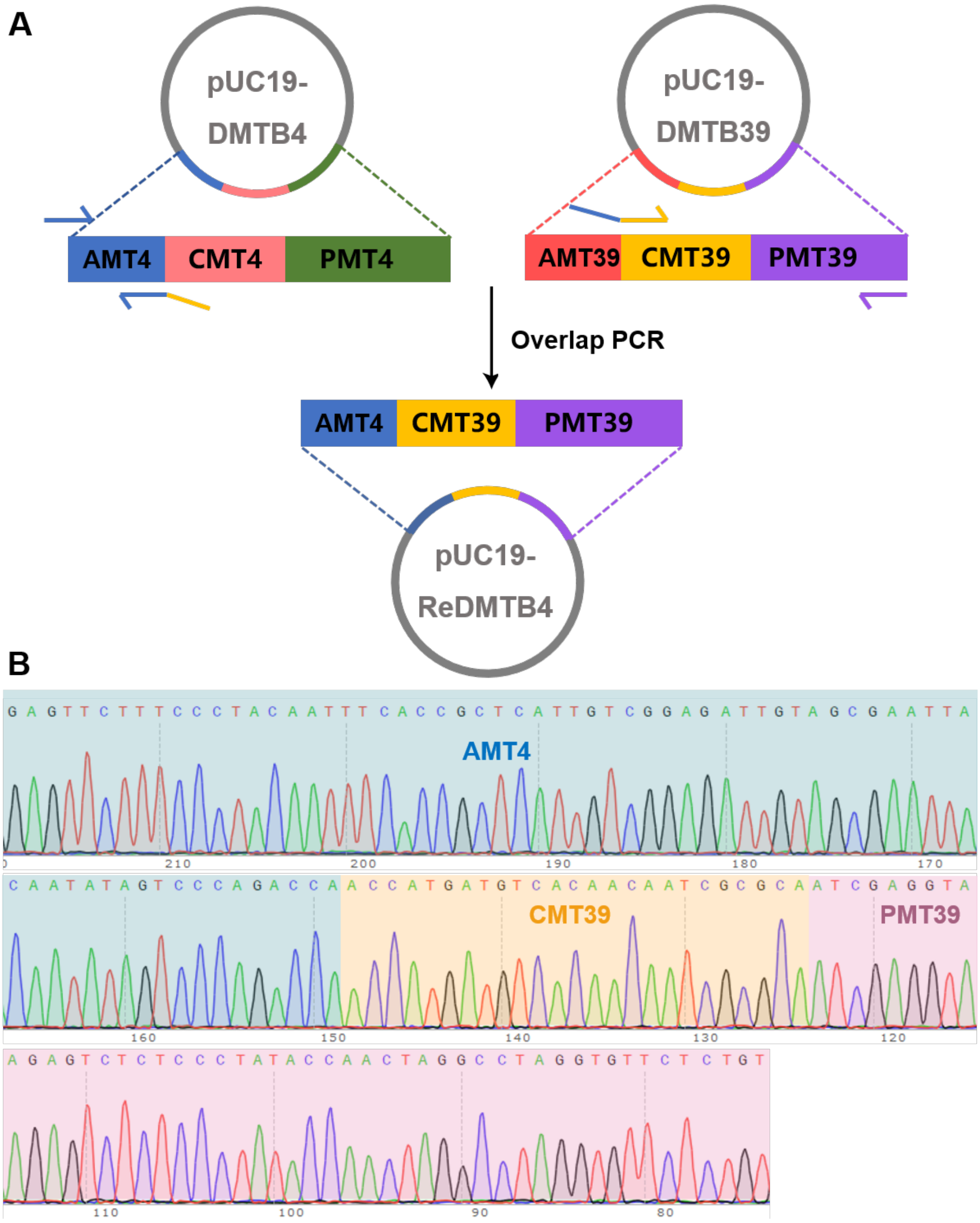


**Fig. S15.** Construction of ReDMTB4. (**A**) ReDMTB4 was constructed by combining the address module (AMT) of DMTB4, which encoded the character “pu”, with the checksum and payload modules (CMT and PMT) of DMTB39, which encoded the character “yin”. This recombinant block retained the positional identity of “pu” while representing “yin”. (**B**) Sequencing analysis of ReDMTB4.

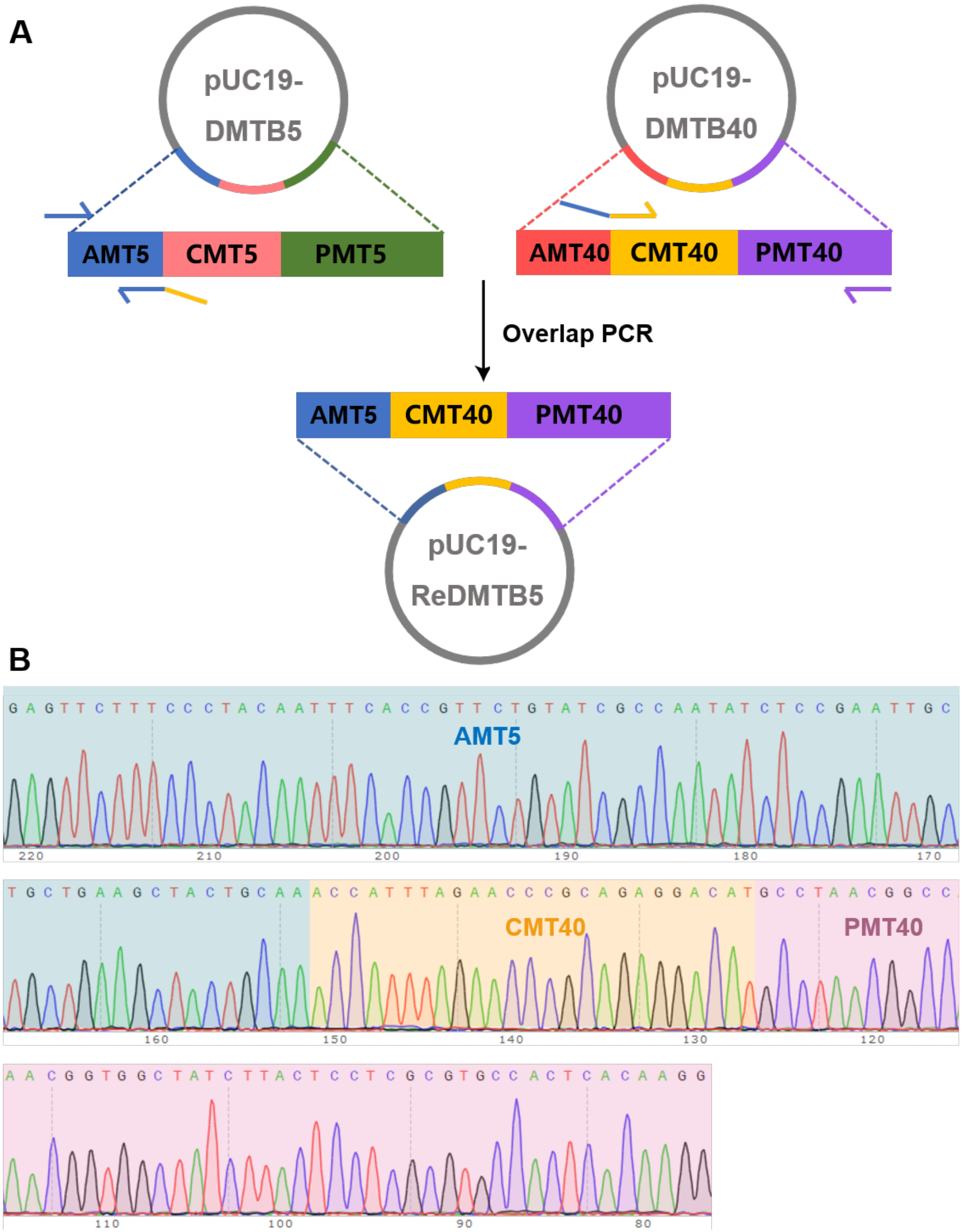


**Fig. S16.** Construction of ReDMTB5. (**A**) ReDMTB5 was constructed by combining the address module (AMT) of DMTB5, which encoded the character “bu”, with the checksum and payload modules (CMT and PMT) of DMTB40, which encoded the character “he”. This recombinant block retained the positional identity of “bu” while representing “he”. (**B**) Sequencing analysis of ReDMTB5.

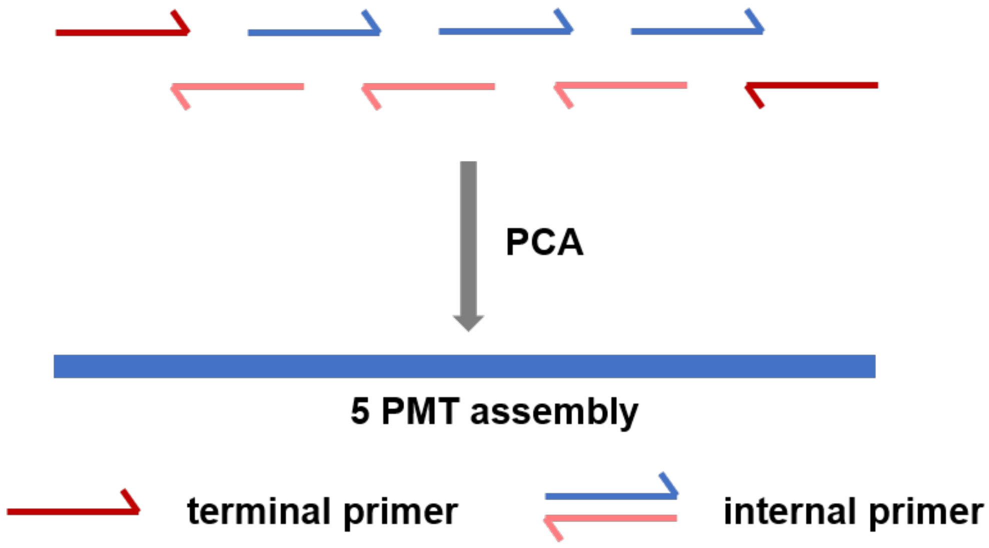


**Fig. S17.** Assembly of five PMTs using PLS DNA polymerase-based PCA.

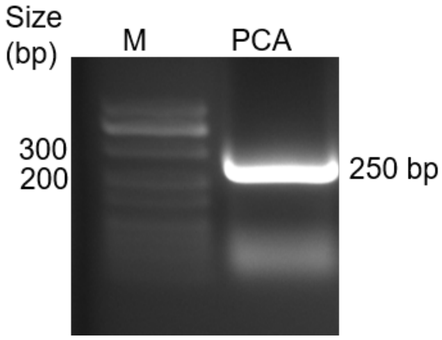


**Fig. S18.** Gel electrophoresis analysis of the PCA products. M: Marker.

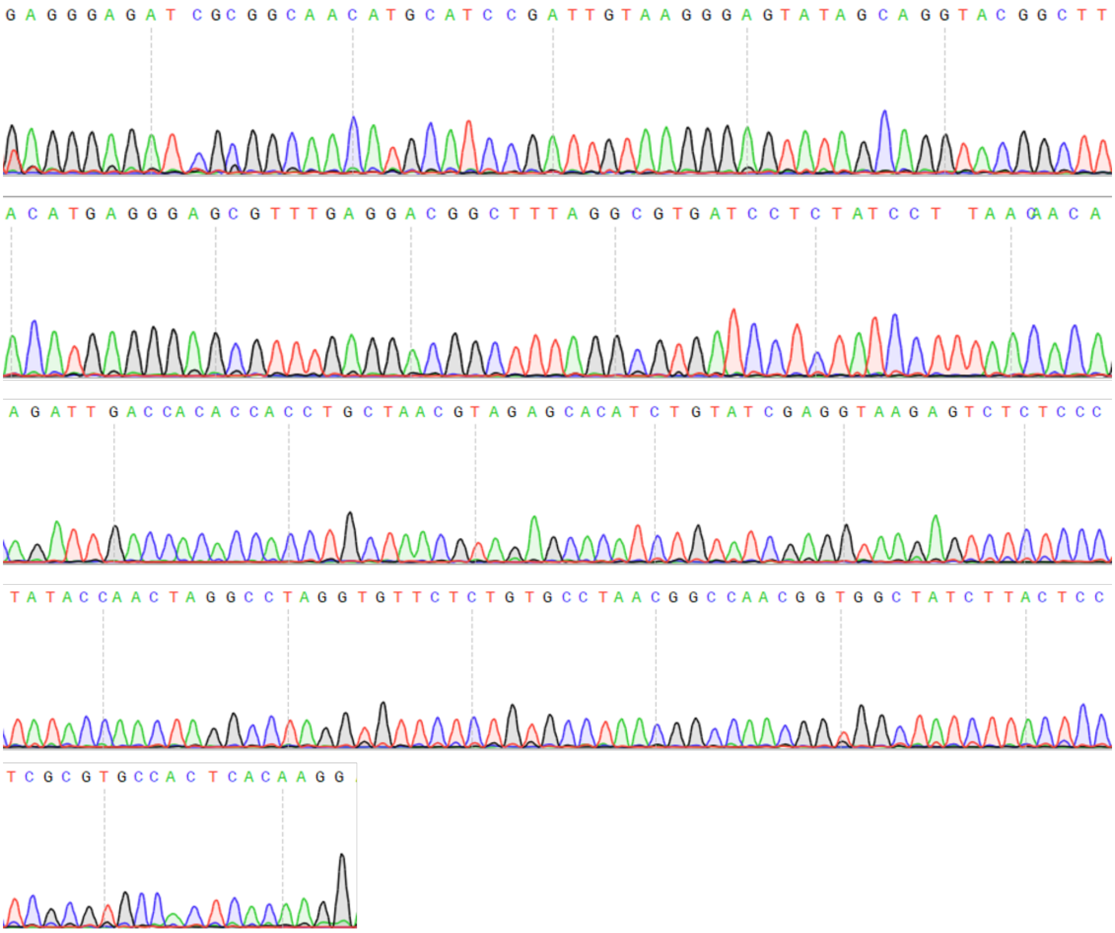


**Fig. S19.** Sequencing analysis of PCA products.

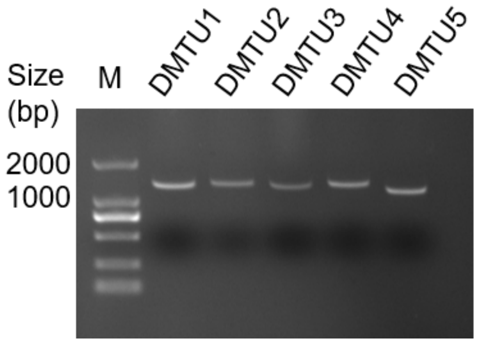


**Fig. S20.** Gel electrophoresis of DMTUs. DMTU1 to DMTU4 each consisted of 1393 bp products, and DMTU5 was 1242 bp in length.

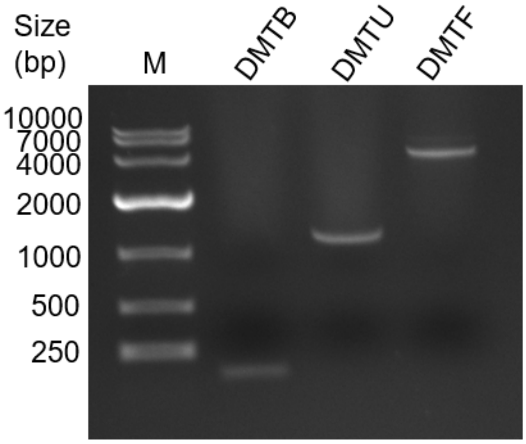


**Fig. S21.** Gel electrophoresis analysis of DMTB1, DMTU4 and DMTF. M: Marker.

**Table S1** DNA sequences of DNA binding proteins.

| DNA binding protein | Sequences (5' to 3') |
|---|---|
| Sso7d | ATGGCGACCGTGAAGTTCAAGTACAAAGGCGAGG AGAAAGAAGTTGATATCAGCAAAATCAAGAAGGT TTGGCGTGTGGGCAAGATGATCTCGTTCACCTATG ATGAGGGCGGCGGTAAGACCGGCAGAGGTGCGGT GAGCGAAAAGGATGCGCCGAAAGAGCTGCTGCAG ATGCTGGAAAAGCAGAAAAAG |
| Sto7d | ATGGTAACAGTTAAATTTAAGTATAAAGGAGAGGA AAAGGAGGTTGATATCTCCAAAATCAAAAAGGTGT GGCGTGTTGGCAAGATGATTTCGTTCACGTACGAC GACAACGGCAAGACCGGTCGCGGTGCGGTGAGCG AAAAAGATGCTCCGAAAGAGTTGCTGCAGATGCT GGAAAAAAGCGGTAAGAAG |
| Sac7d | ATGGTTAAAGTAAAGTTTAAATATAAAGGAGAGGA AAAGGAGGTGGATACCAGCAAAATCAAAAAGGTG TGGCGTGTTGGTAAAATGGTCTCCTTCACGTACGA CGACAACGGCAAAACCGGTCGCGGCGCAGTTAGC GAGAAGGACGCTCCGAAAGAGCTGCTGGATATGTT GGCGCGTGCGGAACGTGAAAAGAAG |

**Table S2** Primer sequences provided in the study.

| Oligo name | Sequences (5' to 3') |
|---|---|
| **2.3 Processivity assay of chimerical DNA polymerases** | |
| M13FAM1-F | FAM-GTTTTCCCAGTCACGACGTTGTAAAACGACGGCC |
| **2.4 Thermal stability of chimerical DNA polymerases** | |
| F | GCAGATTGTACTGAGAGTGCA |
| R | GACGATAGTTACCGGATAAGG |
| **2.5 Salt tolerance assay of chimerical DNA polymerases** | |
| F | GCAGATTGTACTGAGAGTGCA |
| R | GACGATAGTTACCGGATAAGG |
| **2.6 PCR efficiency assay** | |
| λ-1-R (1 kb) | GCAATGGCGATGACGCATCCTCACG |
| λ-2-R (2 kb) | CCATGATTCAGTGTGCCCGTCTGG |
| λ-4-R (4 kb) | CGAAGAGCATCCTCAGGATGTGATGG |
| λ-6-R (6 kb) | GAGATGGCATATTGCTACGCAAGA |
| λ-8-R (8 kb) | GCCTCGTTGCGTTTGTTTGCACG |
| Anchor-λ-F | TCATGCATTGCCTGCTCTGCC |
| **2.11 Random access by DNA polymerases** | |
| JLU-F | TGGCTCATTTCACAATCGGT |
| JLU-R | ATAAATGACCTGCCGTGCAA |
| SKY-F | ATTATTGGCTCCTGCTTGCA |

| | |
|---|---|
| SKY-R | AATGTAGGCGGAAAGTGCAA |
| **2.14 Assembly of DNA movable types via PCA** | |
| PCA-1 | GAGGGAGATCGCGGCAACATGCATCCGATTGTAAGGGAGTATAGCAGGTA |
| PCA-2 | TCCTCAAACGCTCCCTCATGTTAAGCCGTACCTGCTATACTCCCTTACA |
| PCA-3 | CATGAGGGAGCGTTTGAGGACGGCTTTAGGCGTGATCCTCTATCCTTTA |
| PCA-4 | CGTTAGCAGGTGGTGTGGTCAATCTGTGTTAAAGGATAGAGGATCACGC |
| PCA-5 | ACCACACCACCTGCTAACGTAGAGCACATCTGTATCGAGGTAAGAGTCTC |
| PCA-6 | GCACAGAGAACACCTAGGCCTAGTTGGTATAGGGAGAGACTCTTACCTCG |
| PCA-7 | CCTAGGTGTTCTCTGTGCCTAACGGCCAACGGTGGCTATCTTACT |
| PCA-8 | CCTTGTGAGTGGCACGCGAGGAGTAAGATAGCCACCGTTG |

**Table S3.** AMT DNA movable type sequences of Chinese poem.

| No. | Character | AMT | Sequences (5' to 3') |
|---|---|---|---|
| 1 | wang | A1B1C1 | GAGTTCTTTCCCTACAATTTCACCGCTCATTGTCGGAGATTGTAGCGAATTGCTGCTGAAGCTACTGCAA |
| 2 | lu | A1B1C2 | GAGTTCTTTCCCTACAATTTCACCGCTCATTGTCGGAGATTGTAGCGAATTCTCCAGTCAAAAGTCATTA |
| 3 | shan | A1B1C3 | GAGTTCTTTCCCTACAATTTCACCGCTCATTGTCGGAGATTGTAGCGAATGCTATGGCCTTGCGACAGAT |
| 4 | pu | A1B1C4 | GAGTTCTTTCCCTACAATTTCACCGCTCATTGTCGGAGATTGTAGCGAATTACAATATAGTCCCAGACCA |
| 5 | bu | A1B2C1 | GAGTTCTTTCCCTACAATTTCACCGTTCTGTATCGCCAATATCTCCGAATTGCTGCTGAAGCTACTGCAA |
| 6 | /n | A1B2C2 | GAGTTCTTTCCCTACAATTTCACCGTTCTGTATCGCCAATATCTCCGAATTCTCCAGTCAAAAGTCATTA |
| 7 | li | A1B2C3 | GAGTTCTTTCCCTACAATTTCACCGTTCTGTATCGCCAATATCTCCGAATGCTATGGCCTTGCGACAGAT |
| 8 | bai | A1B2C4 | GAGTTCTTTCCCTACAATTTCACCGTTCTGTATCGCCAATATCTCCGAATTACAATATAGTCCCAGACCA |
| 9 | /n | A1B3C1 | GAGTTCTTTCCCTACAATTTCACCGCGTGAGTTCAATCTTATTCCCGAATTGCTGCTGAAGCTACTGCAA |
| 10 | ri | A1B3C2 | GAGTTCTTTCCCTACAATTTCACCGCGTGAGTTCAATCTTATTCCCGAATTCTCCAGTCAAAAGTCATTA |
| 11 | zhao | A1B3C3 | GAGTTCTTTCCCTACAATTTCACCGCGTGAGTTCAATCTTATTCCCGAATGCTATGGCCTTGCGACAGAT |

| | | | |
|---|---|---|---|
| 12 | xiang | A1B3C4 | GAGTTCTTTCCCTACAATTTCACCGCGTGAGTTCAATCTTATTCCCGAATTACAATATAGTCCCAGACCA |
| 13 | lu | A1B4C1 | GAGTTCTTTCCCTACAATTTCACCGTAACGCTAGTCAGGTGAGTCCGAATTGCTGCTGAAGCTACTGCAA |
| 14 | sheng | A1B4C2 | GAGTTCTTTCCCTACAATTTCACCGTAACGCTAGTCAGGTGAGTCCGAATTCTCCAGTCAAAAGTCATTA |
| 15 | zi | A1B4C3 | GAGTTCTTTCCCTACAATTTCACCGTAACGCTAGTCAGGTGAGTCCGAATGCTATGGCCTTGCGACAGAT |
| 16 | yan | A1B4C4 | GAGTTCTTTCCCTACAATTTCACCGTAACGCTAGTCAGGTGAGTCCGAATTACAATATAGTCCCAGACCA |
| 17 | , | A2B1C1 | GTCTTTCGTGAGAGTTCTACCACCGCTCATTGTCGGAGATTGTAGCGAATTGCTGCTGAAGCTACTGCAA |
| 18 | /n | A2B1C2 | GTCTTTCGTGAGAGTTCTACCACCGCTCATTGTCGGAGATTGTAGCGAATTCTCCAGTCAAAAGTCATTA |
| 19 | yao | A2B1C3 | GTCTTTCGTGAGAGTTCTACCACCGCTCATTGTCGGAGATTGTAGCGAATGCTATGGCCTTGCGACAGAT |
| 20 | kan | A2B1C4 | GTCTTTCGTGAGAGTTCTACCACCGCTCATTGTCGGAGATTGTAGCGAATTACAATATAGTCCCAGACCA |
| 21 | pu | A2B2C1 | GTCTTTCGTGAGAGTTCTACCACCGTTCTGTATCGCCAATATCTCCGAATTGCTGCTGAAGCTACTGCAA |
| 22 | bu | A2B2C2 | GTCTTTCGTGAGAGTTCTACCACCGTTCTGTATCGCCAATATCTCCGAATTCTCCAGTCAAAAGTCATTA |
| 23 | gua | A2B2C3 | GTCTTTCGTGAGAGTTCTACCACCGTTCTGTATCGCCAATATCTCCGAATGCTATGGCCTTGCGACAGAT |
| 24 | qian | A2B2C4 | GTCTTTCGTGAGAGTTCTACCACCGTTCTGTATCGCCAATATCTCCGAATTACAATATAGTCCCAGACCA |

| | | | |
|---|---|---|---|
| 25 | chuan | A2B3C1 | GTCTTTCGTGAGAGTTCTACCACCGCGTGAGTTCAATCTTATTCCCGAATTGCTGCTGAAGCTACTGCAA |
| 26 | 。 | A2B3C2 | GTCTTTCGTGAGAGTTCTACCACCGCGTGAGTTCAATCTTATTCCCGAATTCTCCAGTCAAAAGTCATTA |
| 27 | /n | A2B3C3 | GTCTTTCGTGAGAGTTCTACCACCGCGTGAGTTCAATCTTATTCCCGAATGCTATGGCCTTGCGACAGAT |
| 28 | fei | A2B3C4 | GTCTTTCGTGAGAGTTCTACCACCGCGTGAGTTCAATCTTATTCCCGAATTACAATATAGTCCCAGACCA |
| 29 | liu | A2B4C1 | GTCTTTCGTGAGAGTTCTACCACCGTAACGCTAGTCAGGTGAGTCCGAATTGCTGCTGAAGCTACTGCAA |
| 30 | zhi | A2B4C2 | GTCTTTCGTGAGAGTTCTACCACCGTAACGCTAGTCAGGTGAGTCCGAATTCTCCAGTCAAAAGTCATTA |
| 31 | xia | A2B4C3 | GTCTTTCGTGAGAGTTCTACCACCGTAACGCTAGTCAGGTGAGTCCGAATGCTATGGCCTTGCGACAGAT |
| 32 | san | A2B4C4 | GTCTTTCGTGAGAGTTCTACCACCGTAACGCTAGTCAGGTGAGTCCGAATTACAATATAGTCCCAGACCA |
| 33 | qian | A3B1C1 | CTGTGGATGCTGCTACAAACCACCGCTCATTGTCGGAGATTGTAGCGAATTGCTGCTGAAGCTACTGCAA |
| 34 | chi | A3B1C2 | CTGTGGATGCTGCTACAAACCACCGCTCATTGTCGGAGATTGTAGCGAATTCTCCAGTCAAAAGTCATTA |
| 35 | , | A3B1C3 | CTGTGGATGCTGCTACAAACCACCGCTCATTGTCGGAGATTGTAGCGAATGCTATGGCCTTGCGACAGAT |
| 36 | /n | A3B1C4 | CTGTGGATGCTGCTACAAACCACCGCTCATTGTCGGAGATTGTAGCGAATTACAATATAGTCCCAGACCA |
| 37 | yi | A3B2C1 | CTGTGGATGCTGCTACAAACCACCGTTCTGTATCGCCAATATCTCCGAATTGCTGCTGAAGCTACTGCAA |

| | | | |
|---|---|---|---|
| 38 | si | A3B2C2 | CTGTGGATGCTGCTACAAACCACCGTTCTGTATCGCCAATATCTCCGAATTCTCCAGTCAAAAGTCATTA |
| 39 | yin | A3B2C3 | CTGTGGATGCTGCTACAAACCACCGTTCTGTATCGCCAATATCTCCGAATGCTATGGCCTTGCGACAGAT |
| 40 | he | A3B2C4 | CTGTGGATGCTGCTACAAACCACCGTTCTGTATCGCCAATATCTCCGAATTACAATATAGTCCCAGACCA |
| 41 | luo | A3B3C1 | CTGTGGATGCTGCTACAAACCACCGCGTGAGTTCAATCTTATTCCCGAATTGCTGCTGAAGCTACTGCAA |
| 42 | jiu | A3B3C2 | CTGTGGATGCTGCTACAAACCACCGCGTGAGTTCAATCTTATTCCCGAATTCTCCAGTCAAAAGTCATTA |
| 43 | tian | A3B3C3 | CTGTGGATGCTGCTACAAACCACCGCGTGAGTTCAATCTTATTCCCGAATGCTATGGCCTTGCGACAGAT |
| 44 | 。 | A3B3C4 | CTGTGGATGCTGCTACAAACCACCGCGTGAGTTCAATCTTATTCCCGAATTACAATATAGTCCCAGACCA |

**Table S4.** CMT DNA movable type sequences of Chinese poem.

| No. | Character | CMT | Sequences (5' to 3') |
|---|---|---|---|
| 1 | wang | CMT1 | ACCATAGGGTGCGTTTTAGCCAATG |
| 2 | lu | CMT2 | ACCATGACCCTAGATTTGCCATTTG |
| 3 | shan | CMT3 | ACCATAAAAGTTTAAGGTGATTCTC |
| 4 | pu | CMT4 | ACCATCTCATAAACAAGACTTCACA |
| 5 | bu | CMT5 | ACCATAGACCTCTAGTGGCTGATCA |
| 6 | /n | CMT6 | ACCATGATTACTCATTCCGGGTACT |
| 7 | li | CMT7 | ACCATGTGGTAGTCGTTGGAACATG |
| 8 | bai | CMT8 | ACCATATATAGGTGACTGCGATCTA |
| 9 | /n | CMT9 | ACCATATTCCATATTGCTGTCACGT |
| 10 | ri | CMT10 | ACCATTACGAGAGTCGATGACCAAA |
| 11 | zhao | CMT11 | ACCATGAAGCTACAGATCTAAGCCG |

| | | | |
|---|---|---|---|
| 12 | xiang | CMT12 | ACCATCGGTATTTAGCCGGAGTTAT |
| 13 | lu | CMT13 | ACCATTTGGGTGAAAGGTAGTATGT |
| 14 | sheng | CMT14 | ACCATGTGGAACGGAAGTAACTAAT |
| 15 | zi | CMT15 | ACCATGCCTTCCGGGCTAGCAAGTT |
| 16 | yan | CMT16 | ACCATGGGTGCGATGATCGACAGTA |
| 17 | , | CMT17 | ACCATTATTGCTACGGGCGATCTCA |
| 18 | /n | CMT18 | ACCATAACTAATATGCTCAGCCTGA |
| 19 | yao | CMT19 | ACCATGCAGGACACGCTGATATGCA |
| 20 | kan | CMT20 | ACCATACTCTCCGAACCTCGACGGG |
| 21 | pu | CMT21 | ACCATAAAGCTGGCCCTATAGCATG |
| 22 | bu | CMT22 | ACCATAAAGGGACACGGCCTCCAGC |
| 23 | gua | CMT23 | ACCATGGCCGCTATCCACACTGTTA |
| 24 | qian | CMT24 | ACCATGTAGATTATAGTTTGCGCGA |

| | | | |
|---|---|---|---|
| 25 | chuan | CMT25 | ACCATAGGTATATGCTTCTCTGCAC |
| 26 | 。 | CMT26 | ACCATTGTCGCCTCACAACCAGGAC |
| 27 | /n | CMT27 | ACCATGAAACCTCATTTAGCGATAT |
| 28 | fei | CMT28 | ACCATACCCAAGATCATGTGCCAGA |
| 29 | liu | CMT29 | ACCATAGTGCTATAGTTACAGAATG |
| 30 | zhi | CMT30 | ACCATCTAAGTATTGCGATATCTAC |
| 31 | xia | CMT31 | ACCATCCCAATGCGGGAAGTAATGG |
| 32 | san | CMT32 | ACCATAGTACAGGAGAAGGGCCGTG |
| 33 | qian | CMT33 | ACCATGCGCTAGGTTATACTGCTTC |
| 34 | chi | CMT34 | ACCATATTCGAAGAGTTATAACATG |
| 35 | , | CMT35 | ACCATTACGACGACAATGTTAGGAC |
| 36 | /n | CMT36 | ACCATAGAGCCAGAACCACAAACGT |
| 37 | yi | CMT37 | ACCATCTACTGAACAAGGACCACCT |

| | | | |
|---|---|---|---|
| 38 | si | CMT38 | ACCATGGCATCTCCAAGGAATCCTC |
| 39 | yin | CMT39 | ACCATGATGTCACAACAATCGCGCA |
| 40 | he | CMT40 | ACCATTTAGAACCCGCAGAGGACAT |
| 41 | luo | CMT41 | ACCATTGAGTCGTTGGAATCGACGA |
| 42 | jiu | CMT42 | ACCATCGATCTGTAAACCGTCCCAC |
| 43 | tian | CMT43 | ACCATTAAGCAACCCTCAGTAGGTA |
| 44 | 。 | CMT44 | ACCATTATGACCGAGCCTGTACCCG |

**Table S5.** PMT DNA movable type sequences of Chinese poem.

| No. | Character | PMT | Sequences (5’ to 3’) |
|---|---|---|---|
| 1 | wang | PMT1 | TTTCACTGTAGCGGTGTTACGATAGGTTTTAATTCCATGGCTCTACTGTC |
| 2 | lu | PMT2 | TTCTGTTGTTCTCCTTGAGCAATGTTCGTAACGGGTGAGTATGAAGGTTT |
| 3 | shan | PMT3 | TTTGTGTGGTGCAGTATTAGGCTTCCCTATTATTAGCACTACTGCTAGGG |
| 4 | pu | PMT4 | GAGGGAGATCGCGGCAACATGCATCCGATTGTAAGGGAGTATAGCAGGTA |
| 5 | bu | PMT5 | CGGCTTAACATGAGGGAGCGTTTGAGGACGGCTTTAGGCGTGATCCTCTA |
| 6 | /n | PMT6 | TACCAACCAGTAAGTTGAAGCAGTATCAGGACCGACATGAAACAATCTTT |
| 7 | li | PMT7 | TCCCTCACGAGTGGTACGCCGATGAAATCGCGCAAGACGATTAGAACCTA |
| 8 | bai | PMT8 | GGTTCGGGCCTACGGAGGATCAACGTATCAGTAGAGTCTTGCCCTCGCAG |
| 9 | /n | PMT9 | TACCAACCAGTAAGTTGAAGCAGTATCAGGACCGACATGAAACAATCTTT |
| 10 | ri | PMT10 | ATGCTTTGCACTTTAATAGTAGATAATCGGTTGAGGAAGAAGGAACGCTT |
| 11 | zhao | PMT11 | CCAGCCTTTGTCACGTCACAATGCTCTGCAAAGCCGCGGGCGTAGTATGG |
| 12 | xiang | PMT12 | TGTAGTAGCTGACCGTGGACACTGGTTGGAAAATCGTGCAATGTATCGAT |
| 13 | lu | PMT13 | TGGTGTAGTCTCTAGTGATTTGACATATTGAAACATGGTTTTACTACAAA |

| | | | |
|---|---|---|---|
| 14 | sheng | PMT14 | ATAGGTTATATAGTTAAGGGCGGCCATTGTGTAGCGTAGTTGAAGGTGAA |
| 15 | zi | PMT15 | ATCACCCGAACATTGCGATAAAATGAGAGATATTGCCCAACACGAGTATA |
| 16 | yan | PMT16 | ACTTTAGCACGTACATGGGTTGATCAGTGACCCGCCGAGTATTGGCGGCG |
| 17 | , | PMT17 | TGCAATGTCATGGACAAGCACTCCAAAGCATGAATCATTAACGCATAGAT |
| 18 | /n | PMT18 | TACCAACCAGTAAGTTGAAGCAGTATCAGGACCGACATGAAACAATCTTT |
| 19 | yao | PMT19 | CTGCGGCGATCCACAATAGTGCTATTTACGACTCCACTTTCTATTGCTGG |
| 20 | kan | PMT20 | TCAGGTGACACATAGTGCGTCGGCTAGTGATATGAAGATAGCGCCGCGCA |
| 21 | pu | PMT21 | GAGGGAGATCGCGGCAACATGCATCCGATTGTAAGGGAGTATAGCAGGTA |
| 22 | bu | PMT22 | CGGCTTAACATGAGGGAGCGTTTGAGGACGGCTTTAGGCGTGATCCTCTA |
| 23 | gua | PMT23 | GTCATATTGAATCGAGGATCATAGACGCCGCTGCATAGATATCGCCATCC |
| 24 | qian | PMT24 | CTGGTTTGGGCACATATTTATAGCGCGCCTTTAGAGATAGTTAGGCACTA |
| 25 | chuan | PMT25 | AACGCATCGGATGGACGCCTGTAACTGGAAGAAGCCAGGGCAACACGGTA |
| 26 | 。 | PMT26 | TCGTAGTAGTATGCGTTCACTAAAGCTAACGTCCATGGTTTGGGTGACTC |
| 27 | /n | PMT27 | TACCAACCAGTAAGTTGAAGCAGTATCAGGACCGACATGAAACAATCTTT |

| | | | |
|---|---|---|---|
| 28 | fei | PMT28 | ACGCCCAATCTTAGCAACGTAATACGTGAGAAAGTATAGACTGCTGTTTC |
| 29 | liu | PMT29 | GCGGTGGTTATTTGTAACGCAAGAACCTTTCCCATGACTAACATAATATA |
| 30 | zhi | PMT30 | TTTTATGGGCTGAACGCGCAGGAAGCACAGGAATGTTTTGTTTCAGCTCT |
| 31 | xia | PMT31 | CGTTCACCAGAAAAGACGGCTTCCACAACCTGATCCGAGTACCACCCGTC |
| 32 | san | PMT32 | GTATGTTATATGTGCCGATTTATAAAGGATGCGTCAGGCCGCTGTTGAGG |
| 33 | qian | PMT33 | GCAGAAGGGTGGCGGCAAGAATATGAGAGAACAGATAAAGGCGGACATTC |
| 34 | chi | PMT34 | TATGCTGCTGTTCGCGTATGGCGTTCCGCGCGTATTACATCGAGCGCAAT |
| 35 | , | PMT35 | TGCAATGTCATGGACAAGCACTCCAAAGCATGAATCATTAACGCATAGAT |
| 36 | /n | PMT36 | TACCAACCAGTAAGTTGAAGCAGTATCAGGACCGACATGAAACAATCTTT |
| 37 | yi | PMT37 | TGCTGGACAATCGGTTCATAGTACCCACATTAAAGTCCGTGCACTGTTGA |
| 38 | si | PMT38 | TCCTTTAACACAGATTGACCACACCACCTGCTAACGTAGAGCACATCTGT |
| 39 | yin | PMT39 | ATCGAGGTAAGAGTCTCTCCCTATACCAACTAGGCCTAGGTGTTCTCTGT |
| 40 | he | PMT40 | GCCTAACGGCCAACGGTGGCTATCTTACTCCTCGCGTGCCACTCACAAGG |
| 41 | luo | PMT41 | TCAGGAAGGATGTCAAGCTAAGTTGAGCTATCTGGCTTCGAGTTCCAAGT |
| 42 | jiu | PMT42 | AATCGCTTACACTTATATAGTTTGTGGGTTCCCGCCGTTTGGAACCAGAC |

| | | | |
|---|---|---|---|
| 43 | tian | PMT43 | CCAGCCTTAACTGTGGTATTTCCGTACGCAACTCTGCACTACCCAAGAAG |
| 44 | 。 | PMT44 | TCGTAGTAGTATGCGTTCACTAAAGCTAACGTCCATGGTTTGGGTGACTC |

**Table S6.** Error rate analysis of Sanger sequencing during DMTBs assembly.

| | | Number of bases | Identity (%) |
|---|---|---|---|
| Total True | | 50993 | 99.9079 |
| Total error | | 47 | 0.0920 |
| | Insertion (nt) | 2 | 0.0039 |
| | Deletion (nt) | 39 | 0.0764 |
| | Substitution (nt) | 6 | 0.0117 |
| Total bases | | 51040 | |

[9] Bolger AM, Lohse M, Usadel B. Trimmomatic: a flexible trimmer for Illumina

sequence data, Bioinformatics 2014;30:2114-2120.

[10] Vasimuddin M, Misra S, Li H, et al. Efficient architecture-aware acceleration of BWA-MEM for multicore systems, 2019 IEEE International Parallel and Distributed Processing Symposium (IPDPS) 2019;314-324.

[11] Li H. A statistical framework for SNP calling, mutation discovery, association mapping and population genetical parameter estimation from sequencing data, Bioinformatics 2011; 27:2987-2993.
